# The Conserved Transcription Factor Krüppel Regulates the Survival and Neurogenesis of Mushroom Body Neuroblasts in *Drosophila* Adult Brains

**DOI:** 10.1101/2024.06.25.600704

**Authors:** Jin Man, Xian Shu, Haoer Shi, Xue Xia, Yusanjiang Abula, Yuu Kimata

**Affiliations:** School of Life Science and Technology, ShanghaiTech University, Shanghai 201210, China; Department of Bioengineering, School of Engineering and Applied Sciences, University of Pennsylvania, Philadelphia, PA 19104, USA

## Abstract

In various metazoans, including *Drosophila* and humans, neural progenitors exit the cell cycle and are eliminated by the end of development, limiting adult neurogenesis. We demonstrate that in *Drosophila,* the conserved transcription factor *Krüppel* (*Kr*) controls neurogenic capacity of a specific subset of neuroblasts that forms the mushroom body (MBNBs), analogous to the mammalian hippocampus. The *Irregular facet* mutation, which alters *Kr* expression, and neuroblast-specific Kr depletion allow MBNBs, but not other neuroblasts, to persist beyond development and generate neurons in adult brains. Persisting MBNBs express Imp, an RNA-binding protein that promotes neuroblast proliferation and survival. Our results underscore a critical role for Kr in the developmental control of a specific progenitor population, uncovering a novel mechanism controlling adult neurogenesis.

## Introduction

The adult brain is predominantly composed of postmitotic neurons, which lack the capacity for proliferation. This characteristic renders brain tissue, particularly neurons, largely irreplaceable if lost due to damage or degeneration. The loss of neurons is a critical factor in the deterioration of brain function and is associated with various chronic pathologies and organ defects that accompany ageing and degenerative diseases^1,2^. While the brain does contain a minority of proliferative cells, these cells are scarce and predominantly quiescent, further limiting the regenerative potential of the adult brain.

Neural progenitors play a crucial role in brain development, responsible for generating the majority of the brain’s cellular constituents. These progenitors primarily undergo asymmetric cell divisions to self-renew and produce differentiated progeny, which subsequently mature into neurons or glial cells, or partially differentiated intermediate progenitors^3,4^. However, most neural progenitor cells are eliminated during the late stages of development, leaving only a small pool in specific regions of the adult brain, such as the subgranular zone in the hippocampus in mammals. These remaining progenitors are typically slow-proliferating or dormant and can be activated in response to specific physiological or pathological stimuli^5–7^.

*Drosophila* serves as an excellent model organism for studying the development and physiology of the central nervous system, including the behaviour and regulation of neural progenitors. *Drosophila* neural progenitors, neuroblasts (NBs), are highly proliferative and responsible for producing virtually all cells in the fly brain^8,9^. NBs predominantly undergo stereotypical asymmetric cell division to generate a daughter NB and an intermediate progenitor called the ganglion mother cell (GMC), which subsequently divides once to become neurons or glia^3,10^.

The neurogenic and self-renewal capacity of NBs, as well as their cell cycle control and survival, are tightly regulated in a lineage dependent manner during development. At the end of embryogenesis, while many NBs in the abdominal and thoracic segments undergo apoptosis, the majority of NBs in the brain enter a phase of quiescence, only to be reactivated upon larval hatching and food intake^11^. An exception to this regulation are mushroom body neuroblasts (MBNBs), which continue dividing during the embryo-to-larva transition^9^. However, before adult eclosion, all NBs, including MBNBs, are either permanently withdrawn from the cell cycle or eliminated through apoptosis or differentiation^9,12–14^.

Although it was previously believed that no adult neurogenesis occurs in *Drosophila* adult brains, recent studies employing highly sensitive lineage tracing techniques have demonstrated that slow neurogenesis continues beyond adulthood in optic lobes, which can be accelerated upon acute injury^15^. Additionally, quiescent NBs have also been identified in the central brain, where neurogenesis and/or gliogenesis can be induced by physical damage or excessive neuron death^16,17^. Nevertheless, the occurrence of active neurogenesis in the unperturbed adult central brains, as well as the origins and identities of the quiescent NBs, remain unclear.

Temporal transcription factors (tTFs) are a set of transcription factors expressed in NBs and their progeny in temporal cascades during development, which are thought to confer temporal identity to generate the diversity of neural lineages in the central nervous system^18–20^. The temporal sequence and the function of tTFs is particularly well characterised in *Drosophila* embryonic thoracic NB linages. A similar temporal expression pattern of transcription factors is also observed in early brain development in mammals^19,20^. In addition to providing temporal identity, tTFs also regulate the proliferation and survival of NBs during development. In human, misregulation of these TFs may be implicated in childhood cancer^19^.

Krüppel family zinc-finger proteins are an evolutionarily conserved family of transcription factors that play a pivotal role in development, physiology, and stem cell regulation across metazoan species^21,22^. *Drosophila* Krüppel (Kr), a founding member of this family, was first identified as a gap gene during embryonic development^23^. In embryonic brain development, Kr is expressed in many NBs where it acts as a tTF^24,25^. Kr expression has also been reported in neurons during postembryonic stages, suggesting its continued involvement in neural functions^26^. However, the specific roles of Kr in different NB lineages remain largely uncharacterised.

In this study, we elucidate a novel function of the Kr transcription factor in regulating the survival and proliferative capacity of MBNBs in *Drosophila* adult brains. We show that the classic mutation *Irregular facet* (*If*), previously linked to morphological defects in adult eyes^27,28^, deregulates Kr, leading to the unexpected presence of mitotic NBs in the adult central brain. These NBs, identified as MBNBs, appear to bypass the developmentally programmed elimination that occurs during the pupal stage. Our investigations suggest that precise modulation of Kr expression is critical for timing the cell cycle exit and subsequent elimination of MBNBs during and beyond postembryonic development, highlighting a key role for this conserved transcription factor in the longevity and neurogenic capacity of specific neural progenitors. Furthermore, *If* emerges as the first example of a spontaneous mutation that alters adult neurogenesis in metazoans, opening new avenues for understanding the genetic control of neurogenesis in adult brains.

## Results

### The *Drosophila* adult central brain is in a postmitotic state

Recent studies demonstrated that a small fraction of NBs are retained and produce neurons in the optic lobes of the *Drosophila* adult brain and that *de novo* neurogenesis can be induced by physical or physiological stimuli in the central brain^15,17,29^. However, in the unperturbed condition, actively proliferating NBs have never been observed in the central brain. To confirm this, we examined the presence of mitotic cells by immunostaining with an antibody against a mitotic marker, phosphorylated histone H3 Serine 10 (pH3), and found no pH3-positive cells in the control fly adult brains (Fig. 1B). We also performed EdU labelling assays to detect proliferating cells but did not observe any EdU incorporation after 5 days of EdU feeding (Fig. 3A). We also used the Fly-FUCCI system, by which cell cycle phases can be visualised by the expression of the two fluorescent markers that are subject to cell cycle-dependent proteolysis (Fig. 1A)^30^. Virtually all cells in the central brain showed only EGFP-E2F1 1-230 signals but not mRFP-NLS-CycB 1-266 signals, indicating that they are in G1 or G0 phase of the cell cycle (Fig. 1A, B). As an exception, we occasionally observed one or two cells near the antenna lobes in some adult brains that showed EGFP and mRFP signals, but no pH3 signals, indicative of G2 cells (Fig 1B), which may correspond to G2 cells that has been observed in the adult brains during the so-called “critical period”, spanning 5 days from eclosion^7,31^.

**Figure 1.**
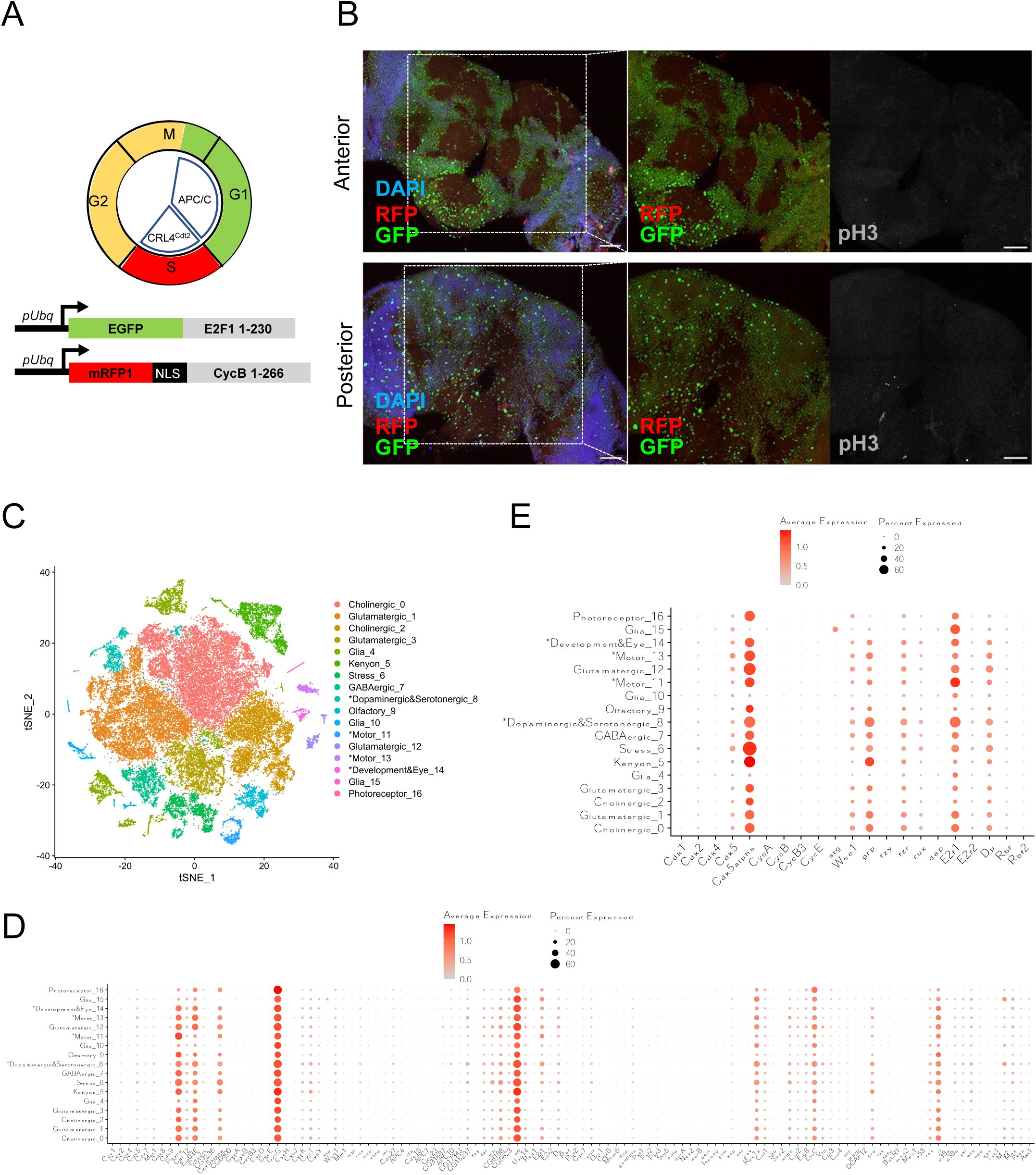
The *Drosophila* adult central brains are post-mitotic. (A) Schematic diagram of the Fly-FUCCI cell cycle reporter. The reporter consists of GFP-tagged E2F11-230 and RFP-tagged Cyclin B1-266, which mark cells in G1/G0 and S/G2/M phases, respectively. (B) Representative confocal images of control *w^1118^* adult central brains ubiquitously expressing Fly-FUCCI reporters (GFP in green, RFP in red), co-stained with DAPI (blue) for DNA and anti-phosphorylated histone H3 serine 10 antibody (pH3, grey). Upper panels show anterior views and lower panels show posterior views. Most cells in the adult central brains express only EGFP-E2F11-230, indicating they are in G1 or G0 phase. Scale bars: 50 µm. (C) tSNE plot showing the clustering of single-cell RNA sequencing data from *Drosophila* adult brains^32^ (Davie *et al.*, 2018). 17 distinct cell clusters are depicted in different colours. An asterisk (*) before cluster names indicates uncertainty in cell type identification. (D) Dot plot showing the expression patterns of 112 CCR genes across the identified 17 cell clusters. The colour intensity of each dot represents the average expression level of the CCR gene, with darker red indicating higher expression levels. The size of each dot indicates the proportion of cells within the cluster expressing the gene, with larger dots representing higher proportions. (E) Detailed view of selected CCR gene expression patterns in the identified cell clusters of the adult brain, extracted from panel D. The expression patterns are displayed as in panel D, emphasising the distribution and intensity of CCR gene expression across different cell types.

To gain insight into the mechanism that controls the postmitotic state of the adult brain, we examined the expression patterns of cell cycle regulator (CCR) genes using recent single-cell RNA sequencing (scRNAseq) data of the *Drosophila* adult brain^32^. The original data were obtained and cells were organised into 17 clusters based on their gene expression profiles (Fig. 1C). The transcription levels of 112 selected CCRs were then analysed in each cell cluster (Fig. 1D, E, S1). Most positive CCRs, typically associated with promoting cell cycle transitions, such as cyclins, cyclin-dependent kinases (Cdks), mitotic kinases, and components of the replication machinery, are minimally expressed across all neuronal and glial cell clusters (Fig. 1D, E, S1). In contrast, some negative CCRs, which function to delay or block cell cycle progression and/or inhibit the activity of positive CCRs, including *Wee1*, the *Drosophila* FZR1 orthologue *fizzy-related* (*fzr*), a CDK inhibitor *roughex* (*rux*), the DNA licensing inhibitor *geminin* and the checkpoint kinase *grape* (*grp*, the CHK1 orthologue), were expressed at relatively high levels in many clusters (Fig. 1D, E, S1). A comparable analysis using the scRNAseq data of larval brains^33^ showed that while a few negative CCRs such as *fzr* are more pronounced in neuronal clusters, the expression of most CCRs associated with cell cycle suppression remains constant across both neuroblast and neuronal clusters (Fig. S2). This result suggests that repression of positive CCRs and sustained expression of negative CCRs may cooperate to maintain the postmitotic state of the adult brain.

However, there were some exceptions: certain positive CCRs, E2F1, the transcription activator that drives G1/S transition, and its co-activator Dp, were highly expressed across various neuronal and glial clusters as in NBs (Fig. 1D, E, S1, S2). Given the known functions of mammalian E2F1 in regulating migration and apoptosis in postmitotic neurons^34–36^, it is plausible that E2F1 may also serve neuron-specific functions in *Drosophila* beyond cell cycle control. Additionally, some non-canonical CDKs, Cdk5, Cdk5α, Eip63E/CDK10, Pitslre/Cdk11 and Cdk12, and their known partner cyclins, CycG, CycK, CycT and CycY, showed constitutively high expression patterns or upregulation in some neuronal cell clusters (Fig. 1D, S1, S2). These CDKs have been reported to regulate transcription or neuronal activity in addition to cell cycle regulation^37,38^. Such non-cell cycle functions may be involved in specialised neuronal functions.

Moreover, several CCRs that act through ubiquitin-dependent protein degradation, such as E2 enzymes, *effete* (*eff,* the Ube2d2 orthologue), *vihar* (*vih*, the Ube2C orthologue) and *CG8188* (the Ube2S orthologue), and some components of APC/C and SCF E3 complexes, exhibit modest to high expression in neural clusters (Fig. 1D, S1, S2), highlighting the critical role of the ubiquitin-proteasome pathway in neurons^39^.

We also performed more detailed analysis on CCR expression in the larger number of cell clusters, 84 clusters, which were identified applying the clustering and annotation methods used in Davie *et al.*^32^, and observed largely similar expression patterns (Fig. S3).

In summary, our transcriptome analysis suggests that a majority of CCRs are likely to regulate the postmitotic state of the adult brains through their canonical functions while a subset of CCRs, such as E2F1, may regulate specific functions of neurons and/or glia independently of cell cycle regulation.

### The presence of mitotic cells in the adult central brain of the *Irregular facet* mutant

During the analysis of cell cycle states in adult brains, an unexpected phenotype was observed in progeny of one of our laboratory stocks, *w^1118^; If/Cyo; mb247-gal4, UAS-mCD8::GFP* (*If/CyO; mb247>mCD8::GFP*). Among the progeny from the crosses between this line and either *w^1118^* or an *E2F1RNAi* line (*v100900* from the Vienna Drosophila Resource Center, VDRC), we observed a few cells in each hemisphere showing clear pH3 signals in the dorsoposterior parts of the adult central brains (Fig. 2A-C, Table 1). These cells exhibited highly condensed chromosomes decorated with pH3 signals, resembling mitotic chromosomes. Intriguingly, we were able to detect these potential mitotic cells in adult brains as long as 14 days after adult eclosion, much beyond the critical period (Table 1)^7,31^. This observation was quite surprising, given the results from our analysis with wild type adult brains (Fig. 1, 3A) and the current consensus that no cell proliferation occurs in the *Drosophila* adult central brain in an unperturbed condition^2,7,40^.

**Figure 2.**
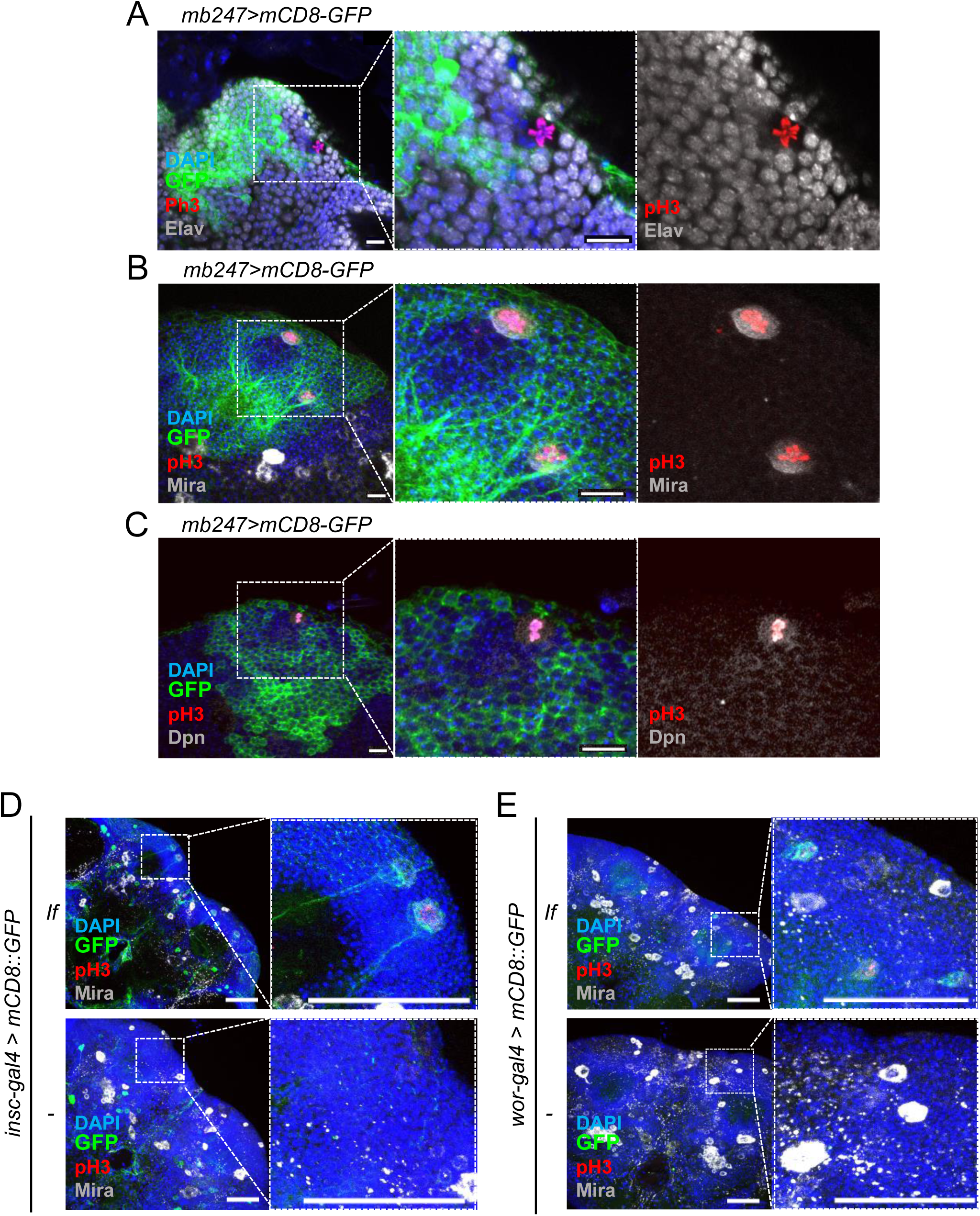
Mitotic neuroblasts in the adult central brain of the *Irregular facet* mutant. (A-C) *If* mutant adult brains expressing mCD8::GFP driven by a mushroom body (MB)-specific driver *mb247*-gal4 (green) were dissected and stained with DAPI (blue), the mitotic cell marker phosphorylated histone H3 serine 10 (pH3, red), and either the pan-neuron marker Elav, neuroblast (NB) marker Miranda (Mira), or Deadpan (Dpn, grey). Mitotic cells with chromosomal pH3 signals were observed in the central brains of If mutant adult flies. These cells did not express Elav (A) but did express Mira (B) and Dpn (C). Mira exhibited a crescent-like asymmetric cortical localization pattern, characteristic of mitotic NBs (B). Scale bars: 10 µm. (D, E) *If* mutant (*If,* top panels) and control (−, lower panels) adult brains expressing mCD8::GFP (green) driven by NB-specific drivers *insc-*gal4 (D) and *wor*-Gal4 (E) were stained with DAPI (blue), anti-pH3 antibody (red), and anti-Mira antibody (grey). Mitotic and interphase NBs, indicated by co-expression of Mira and GFP, were detected in the dorsoposterior regions of the central brains in *If* mutants but not in control. Scale bars: 50 µm.

**Table 1.** Adult central brains with pH3-positive cells in *If/CyO; mb247>mCD8::GFP* progeny at different ages.

| Age<br>(After eclosion) | crossed with *w<sup>1118</sup>* |  |  | crossed with *E2F2 RNAi* |  |  |
| --- | --- | --- | --- | --- | --- | --- |
|  | pH3<sup>+</sup> | Total | Ratio | pH3<sup>+</sup> | Total | Ratio |
| 1 day | 4 | 15 | 26.7% | 18 | 58 | 31.0% |
| 3 days | 9 | 31 | 29.0% | 9 | 41 | 22.0% |
| 7 days | 9 | 40 | 22.5% | 12 | 51 | 23.5% |
| 14 days | 8 | 39 | 20.5% | 6 | 37 | 16.2% |

Number of adult brains with pH3 signals and total number analysed.

To identify the cause of the phenotype, we first carefully examined the experimental condition and genetic components of the parental *If/CyO; mb247>mCD8::GFP* line. Since the above experiment was conducted at 29°C, a suboptimal temperature for fly growth, the phenotype might potentially be caused by heat stress. To test this possibility, we grew *If/CyO; mb247>mCD8::GFP* flies and control flies, *w^1118^*, at three different temperatures, 29°C, 25°C, the optimal temperature for fly raising, and 19°C, a temperature commonly used for long-term maintenance of fly stocks. While no mitotic cells were observed in adult brains of the control at any of the temperature, *If/CyO; mb247>mCD8::GFP* adult brains showed mitotic cells at all the three temperatures, although somewhat higher penetrance of the phenotype was observed at higher temperature (Table 2). Thus, incubation temperature appears not to be responsible for the adult brain phenotype.

**Table 2.**
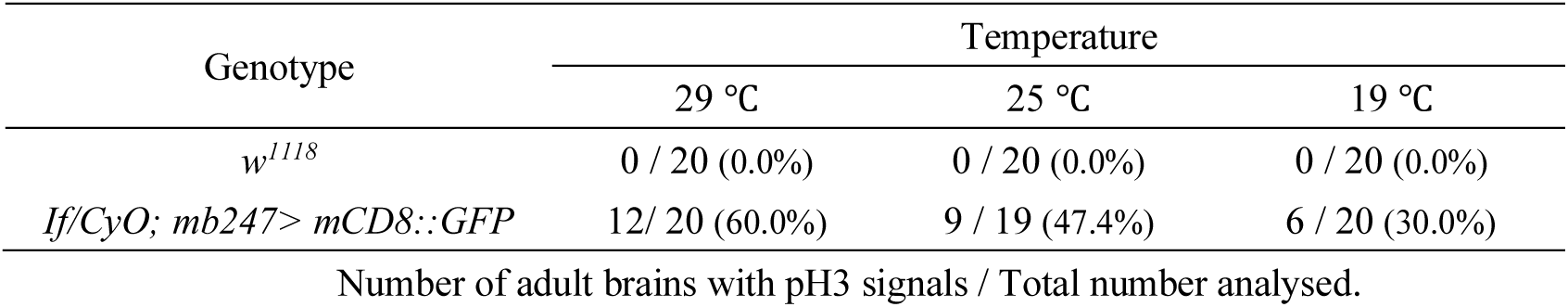
Adult brains with ectopic mitotic cells at different temperatures.

| Genotype | Temperature |  |  |
| --- | --- | --- | --- |
|  | 29 °C | 25 °C | 19 °C |
| *w<sup>1118</sup>* | 0 / 20 (0.0%) | 0 / 20 (0.0%) | 0 / 20 (0.0%) |
| *If/CyO*; *mb247>mCD8::GFP* | 12 / 20 (60.0%) | 9 / 19 (47.4%) | 6 / 20 (30.0%) |

Number of adult brains with pH3 signals / Total number analysed.

We next asked if a certain genetic component of the *If/CyO; mb247>mCD8::GFP* line may induce the adult brain phenotype. We first analysed genetic components on the third chromosomes: a mushroom body neuron-specific driver, *mb247-*gal4^41^ and a fluorescent reporter *UAS-mCD8::GFP*^42^. Neither adult flies carrying *mb247-gal4* nor *UAS-mCD8::GFP* showed any mitotic cells in central brains. We next assessed the involvement of genetic elements in the second chromosomes, the *CyO* balancer chromosome and the second chromosome carrying a dominant mutation *Irregular facet* (*If*)^43^. We conducted co-segregation analysis taking advantage of their dominant phenotypes of *If* and *Curly* (*Cy*, on CyO)^44^, visible in adult flies (narrowed and rough adult eyes and curly wings, respectively). We crossed the *If/CyO; mb247>mCD8::GFP* line with the *w^1118^* line and analysed the adult brains of their progeny. We observed that many progeny that inherited the *If-*bearing second chromosome, but none of the progeny that inherited *CyO*, showed mitotic cells in adult brains (Table 3). We also performed co-segregation analysis using the *w^1118^; If/Cyo; Sb/TM6B* line, from which the *If/CyO; mb247>mCD8::GFP* line derived, and observed co-segregation of *If* and the adult brain phenotype (Table 3). We also detected mitotic cells in the adult brains of another *If* mutant stock that was obtained from Shanghai Fly Center (*w^1118^; If/CyO,* BCF360) (6 in 15 brains, 40.0%). Furthermore, to examine a possible involvement of additional genetic elements in the *If-*bearing second chromosome in this phenotype, we allowed recombination between the *If-*bearing chromosome with the wild type chromosome to assess the co-segregation of the *If* mutation and the adult brain phenotype. While none of non-*If* progeny showed mitotic cells, the majority of *If* recombinants possesses dividing cells (Table 4). Together, these results strongly suggest that the *If* mutation is the cause for the appearance of mitotic cells in the adult central brain.

**Table 3.** Number of flies detected the proliferating cells in adult brain with *If* or *Cyo*.

| Crosses | Temperature | Phenotype of adult progeny |  |
| --- | --- | --- | --- |
|  |  | *If* | *Cy* |
| *w<sup>1118</sup>; If/CyO; mb247-gal4, UAS- mCD8::GFP* | 29 °C | 14/22 (63.6%) | 0/22 (0.0%) |
| *x w<sup>1118</sup>* | 25 °C | 3/20 (15.0%) | 0/18 (0.0%) |
| *w<sup>1118</sup>; If/CyO; Sb/TM6B* | 29 °C | 17/19 (89.5%) | 0/17 (0.0%) |
| *x w<sup>1118</sup>* | 25 °C | 17/29 (58.6%) | 0/22 (0.0%) |

Number of adult brains with pH3 signals / Total number.

**Table 4.** Co-segregation of *If* and the adult brain phenotype in *If* mutant recombinants.

| Recombinant genotype | *If*/+ | +/+ |
| --- | --- | --- |
| **Adult brains with EdU+ clones** | 18/26 (69.2%) | 0/24 (0.0%) |

### The *If* mutation causes the retention of proliferative Mushroom body neuroblasts in the adult central brain

We next wished to determine the identity of the mitotic cells in *If* mutant adult brains. These mitotic cells are not neurons: they are much larger in size compared to their surrounding neurons and do not express the pan-neuronal marker Elav (Fig. 2A). All the mitotic cells in *If* mutant adult brains were expressing the NB markers Miranda (Mira) and Deadpan (Dpn, Fig. 2B, C) and also showed mCD8::GFP induced by NB-specific drivers, *insc-*gal4^45^ and *wor-* gal4^46^ (Fig. 2D, E). These data indicate that the mitotic cells present in the *If* mutant adult brains are NBs. Notably, these NBs exhibite a “crescent”-like, asymmetric localisation of Mira at the cell cortex during mitosis and are surrounded by smaller cells not expressing NB markers (Fig. 2B-E). Thus, similar to conventional NBs, these NBs likely undergo asymmetric cell division to produce a daughter NB and a differentiating daughter cell.

It is noteworthy that while interphase NBs, labelled by Mira and *insc* or *wor>*mCD8::GFP but not pH3, were also observed in the dorsal posterior region of *If* mutant adult brains, neither interphase nor mitotic NBs were detectable in the comparable region of the control adult brains (Fig. 2D, E). This suggests that the *If* mutation likely causes either the retention or the *de novo* formation of NBs, instead of the reactivation of quiescent NBs that may also exist in the wildtype adult brain. We also noted that several cells labelled by either Mira, *insc* or *wor>*mCD8::GFP were scattered throughout the adult central brain of both wild type and the *If* mutant (Fig. 2B, D, E). However, these cells did not co-express these NB markers, making their identities unclear.

We further characterised these anomalous NBs in *If* mutant adult brains by labelling their cell clones using 5-Ethynyl-2’-deoxyuridine (EdU) incorporation. EdU incorporation was observed in NB clones located on the dorsoposterior surface above the mushroom body, with one to four clones in each hemisphere (Fig. 3A, 4C, S5B), a number comparable to that of the mushroom body NBs (MBNBs)^9^. It has been shown that MBNBs are eliminated through apoptosis at the late pupal stage before the end of development^9,47,48^. We hypothesised that the anomalous NBs in *If* mutant brains may be MBNBs that persist into adulthood, extending beyond their usual developmental timeframe.

**Figure 3.**
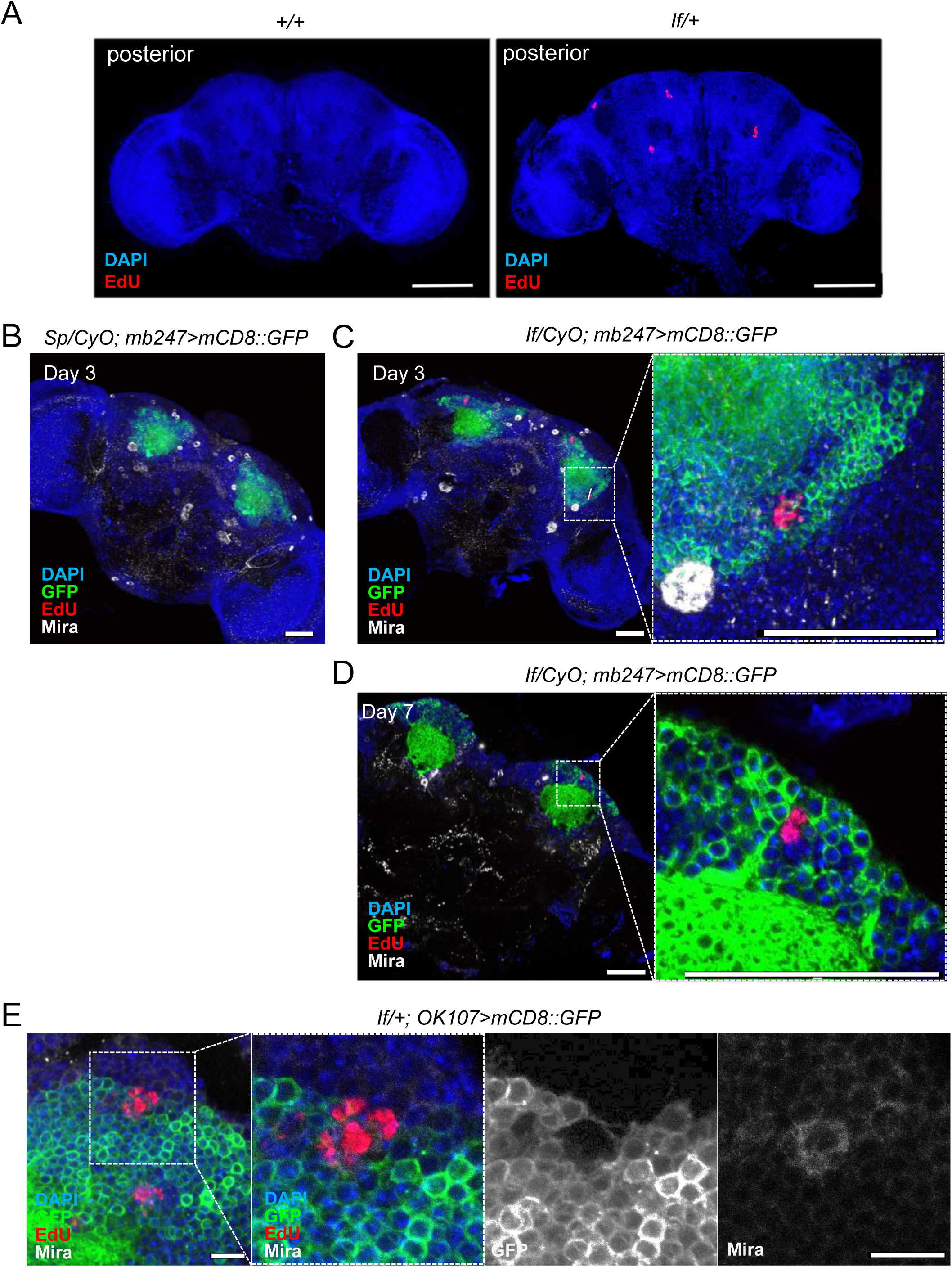
Mushroom body neuroblasts continue proliferating and producing neurons in the *If* mutant adult central brain. (A) EdU labelling was performed in 3 to 5-day-old wild type (*w^1118^; +/+)* and *If* mutant (*w^1118^; If/+*) adult flies and brains were dissected for staining with DAPI (blue) and EdU(red). EdU incorporation was observed in up to 4 clones per hemisphere in *If* mutants. Scale bars: 100 µm. (B-D) EdU labelling was performed in in 3 to 5-day-old control (*Sp/CyO; mb247>mCD8::GFP*) and *If* mutant (*If/CyO; mb247>mCD8::GFP*) adult flies. Representative images of adult brains after 3 days (B, C) or 7 days (D) of a chase after EdU removal. In *If* mutant brains, EdU-labelled cells (red) were incorporated into the existing MBs marked by mCD8::GFP driven by *mb247-* gal4. Scale bars: 50 µm. (E) Representative images of the *If* mutant brains expressing mCD8::GFP in MB lineages using *OK107*-gal4 after EdU labelling. EdU incorporation was observed in MBNBs, marked by co-expression of GFP and Mira. Scale bars:10 µm.

To test this hypothesis, we traced the fate of the progeny of the dividing NBs using EdU labelling chase experiments. Young adult flies of the *If* mutant (*If/CyO; mb247>mCD8::GFP*) and control (*Sp/CyO; mb247>mCD8::GFP*), which harbours another dominant mutation *Sp*, instead of *If*, on the second chromosome, were fed with EdU-containing food for three days, allowing its incorporation into the DNA of all the cells produced during this labelling period. After EdU removal, the flies were incubated for an additional three or seven days, and their brains were then dissected for EdU signal detection. In all *If* mutant brains, EdU incorporation was observed in NB clones in each hemisphere, whereas none of the control brains showed EdU-positive clones (Fig. 3B-D). The EdU-positive clones, composed of up to approximately 10 cells, were embedded in the cell bodies of MB neurons, marked by *mb247>*mCD8::GFP (Fig. 3C, D).

We also employed *OK107-*gal4, the driver that expresses Gal4 in MBNB lineages including MBNBs themselves^49^, with EdU labelling and found that some EdU-positive clones in *If* mutant brains contained cells co-expressing *OK107*-gal4-induced mCD8::GFP and Mira, indicating their MBNB identity (Fig.3E). These data have demonstrated that the active NBs present in the *If* mutant adult brain are MBNBs capable of producing neurons that integrate into the adult MBs, which are likely to have escaped their elimination during pupal stages.

Notably, despite the prolonged neurogenesis, we did not observe significant alterations in the size and morphology of the MBs in *If* mutant brains compared to the control (Fig. S4). The number of additional neurons produced in *If* mutant adult brains appears insufficient to impact the overall structure of the MBs.

### The conserved transcription factor *Krüppel* regulates the survival and proliferation capacity of the MBNB

We next investigated the mechanisms by which the *If* mutation leads to the retention and prolonged neurogenesis of MBNBs in the adult brain. *If* was initially identified as a dominant mutation that results in severe morphological defects in the adult eye^27^ Subsequent studies suggest that *If* is a gain-of-function mutation in the *Krüppel* (*Kr*) gene, causing its aberrant expression. Specifically, *Kr* was found to be highly expressed in the differentiated region of the eye imaginal disc, which likely accounts for the *If* mutant eye phenotype^27,28^. While its effect on eye development is well documented, potential impacts of the *If* mutation on other tissues, including the brain, have not been explored.

We hypothesised that in *If* mutants, *Kr* misexpression in MBNBs affects their normal developmental control, leading to their persistence into the adult brain. To test this hypothesis, we specifically reduced *Kr* expression through RNA interference (RNAi) in NBs in *If* mutants. The induction of Kr-targeted short hairpin RNA (*Kr*shRNA, hereinafter) using the NB-specific *insc-*gal4 driver partially suppressed the formation of mitotic MBNBs in *If* mutant adult brains, as indicated by a reduced number of mitotic NBs and EdU-positive clones (Fig. S5). However, to our surprise, when *Kr* RNAi was induced using another RNAi line composed of a double-strand RNA targeting *Kr* (v104150 from the VDRC. *KrdsRNA#1*, hereinafter), we did not observe any rescue of the adult brain phenotype in the *If* mutant (Fig. S5). Intriguingly, the NB-specific induction of *KrdsRNA#1*, but not *Kr*shRNA, in the wild-type background induced a phenotype that was comparable to the phenotype caused by the *If* mutation (Fig. 4, S5B). Similar to *If* mutants, *insc>KrdsRNA#1* adult files retained actively dividing MBNBs beyond 21 days after eclosion (Fig. 4C). The same phenotype was also observed when another *Kr* dsRNA line (v40871 from VDRC. *KrdsRNA#2*, hereafter) was used (Fig. 4B, C), ruling out the possibility of an unexpected effect of the *KrdsRNA#1* construct on NB proliferation. The differences observed between the *KrdsRNA* lines and *KrshRNA* line can be attributed to the higher efficiency of the dsRNA lines in Kr depletion compared to the shRNA line: when induced in larval eye imaginal discs of *If* mutants using *eyeless-*gal4 driver (*ey-*gal4), the *Kr*dsRNA constructs more fully depleted Kr proteins that were ectopically expressed in the differentiated region of eye imaginal discs of *If* mutants (Fig. S6A, B) and more efficiently restored the morphological defects of the adult eyes than the *KrshRNA* line (Fig. S6C). Together, these results suggest that the endogenous level of Kr in NBs is critical for the timely elimination of MBNBs before the end of development.

**Figure 4.**
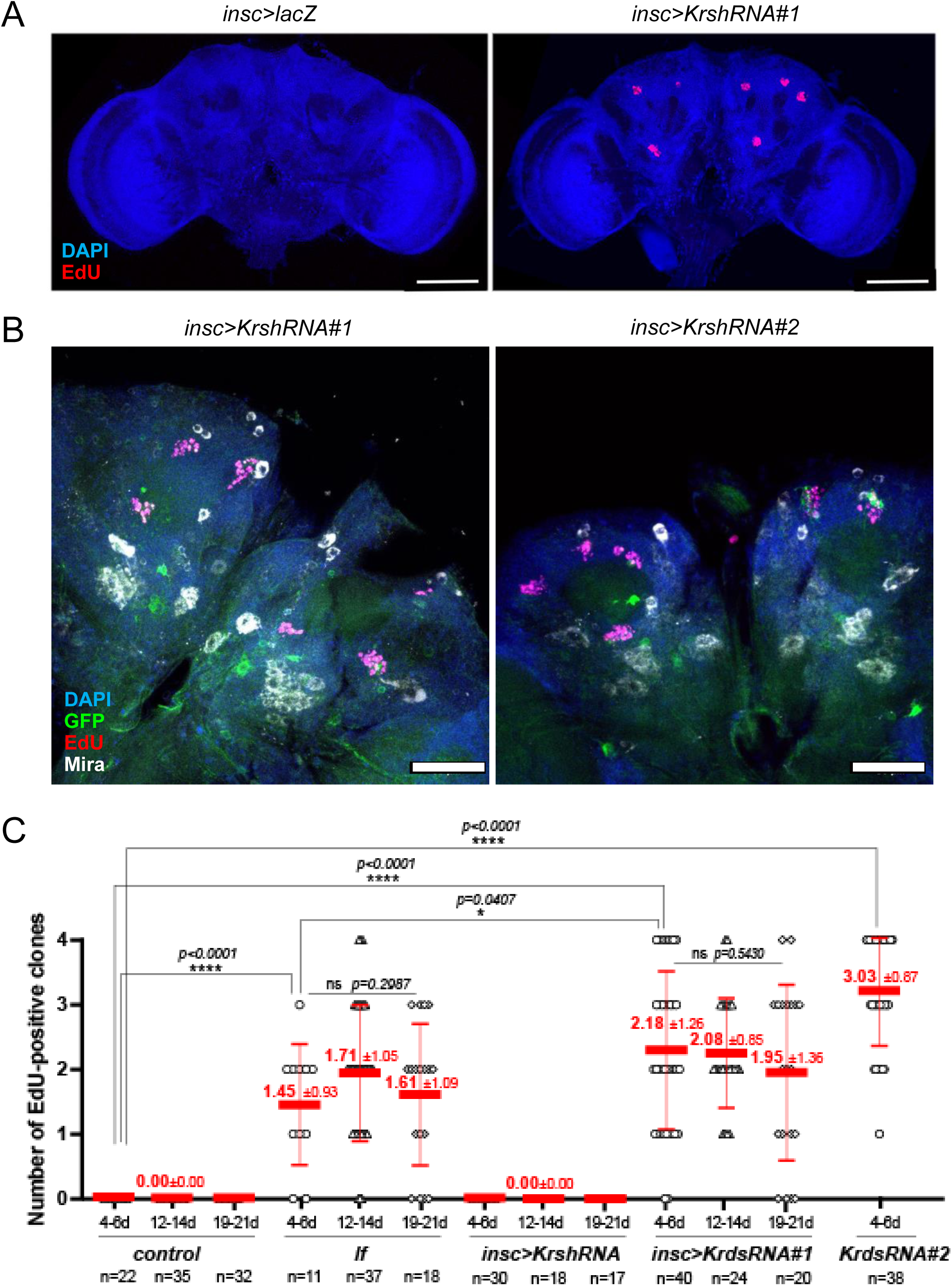
Full depletion of Kr by KrdsRNA in NBs causes MBNB extension and prolonged neurogenesis. (A) EdU labelling was performed in 3 to 5-day-old adult flies of the control (*insc>lacZ*) and those inducing *KrdsRNA* in NBs (*insc>KrdsRNA#1*). Adult brains were dissected for staining with DAPI (blue) and EdU(red). EdU incorporation was observed in *insc>KrdsRNA#1* adult brains. Scale bars: 100 µm. (B) Representative images of adult central brains from *insc>KrdsRNA#1* (left) and *insc>KrdsRNA#2* (right). DNA is shown in blue, *insc*-gal4-driven mCD8::GFP in green, and EdU in red. Both *insc>KrdsRNA#1* and *insc>KrdsRNA#2* exhibited EdU-positive MBNB clones in a wild type background. Scale bars: 50 µm. (C) EdU labelling was performed on adult flies of the specified strains aged 4 to 6 days, 12 to 14 days, and 19 to 21 days after eclosion. Brains were dissected, and EdU-positive cell clones per hemisphere were quantified and displayed in scatter dot plots. The means and standard deviations are represented by thick and thin red bars, respectively, with actual values annotated above in red. ‘n’ indicates the number of brains analysed in each condition. Statistical significance was determined using the Mann-Whitney U test, with significance levels denoted as: * (p ≤ 0.05), ** (p ≤ 0.01), *** (p ≤ 0.001), **** (p < 0.0001). ‘ns’ indicates non-significant results.

We also tested if overexpression of *Kr* in NBs can mimic the *If* mutant phenotype in wild-type adult brains, utilising the *UAS-Kr::V5* transgene whose functionality has been validated^50^. However, the induction of Kr::V5 using NB-specific drivers, *insc-*gal4*, wor-*gal4 or *Elav-*gal4, all led to lethality during early larval stages. When *Kr::V5* expression was induced after the embryonic stage using temperature-sensitive Gal80 (*pTUB-gal80ts*^51^), the flies were able to develop adult. Yet, no MBNBs were observed in the adult brains (Table 5). In addition, our analysis of Kr expression using Kr-specific antibodies did not observe higher expression of Kr in MBNBs in developing or adult brains of *If* mutants, compared to controls, using Kr-specific antibodies (Fig. 5, S7, S8). These data suggests that the *If* mutation may not cause MBNB retention simply by overexpressing *Kr* in MBNBs. Although the *If* mutation has been mapped close to the *Kr* gene^27^, its exact mutation site and its effect on *Kr* expression outside eye imaginal discs have not been characterised. Due to this uncertainty, in our subsequent analysis, we focused on the role of endogenous *Kr* in MBNB regulation using the *KrdsRNA* lines.

**Figure 5.**
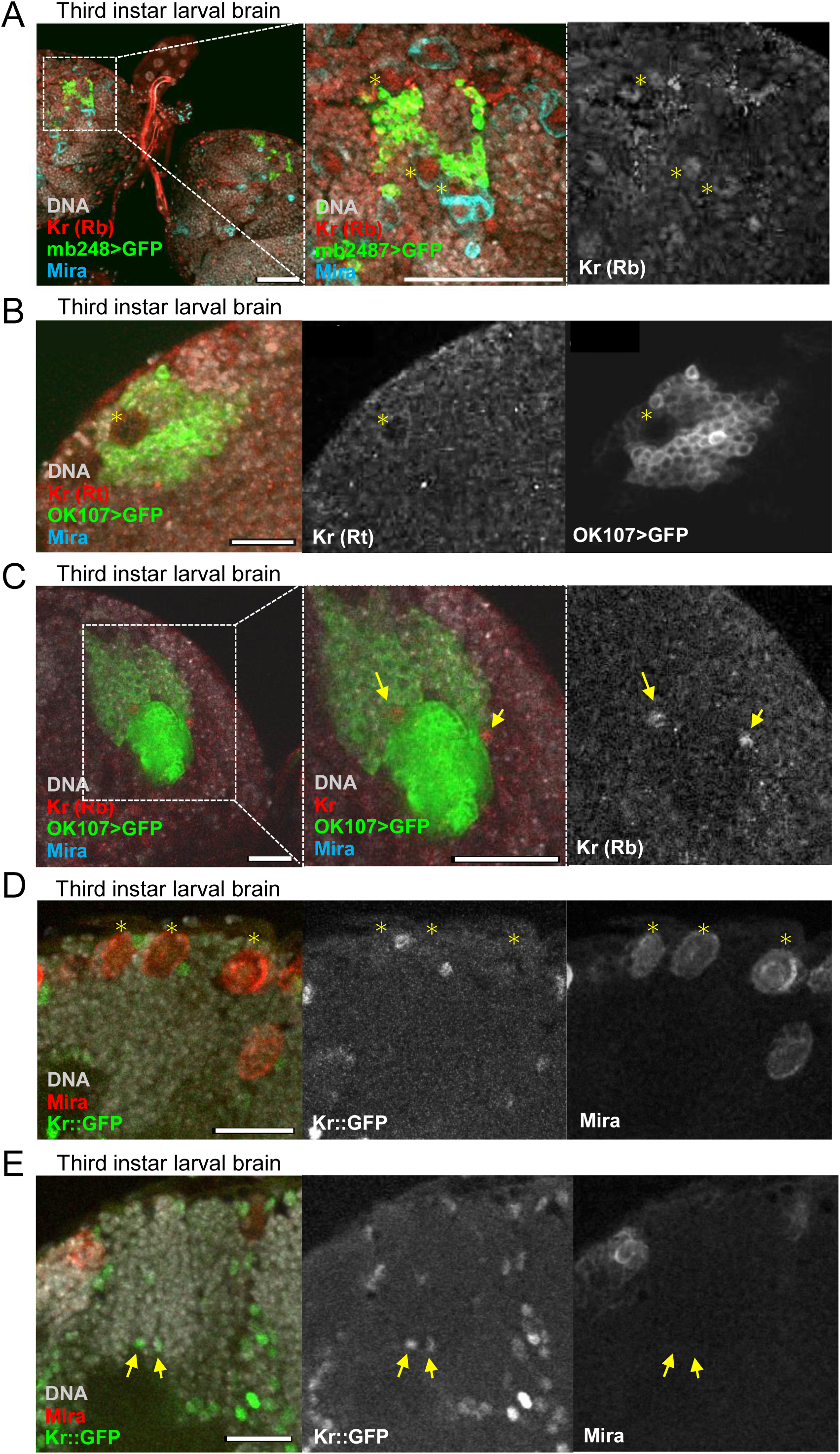
Kr is expressed in MBNBs in larval brains. (A-C) Representative images from the dorsoposterior region of wild-type third instar larval brains, highlighting MB cell bodies and the Calyx. Kr proteins were visualised using Kr-specific antibodies^56^. In merged images, Kr signals are shown in red, DNA in grey, *OK107*>GFP in green, and Mira in cyan, with accompanying mono-coloured views for each signal in grey. (A, B) Magnified views feature MBNBs identified by the co-localisation of *OK107*>GFP and Mira (marked by asterisks), showing weak Kr signals detected by a Kr antibody raised in a rabbit (Kr (Rb), A)^56^ and in a rat (Kr (Rt), B)^57^ (C) Arrows point MBNB lineage cells near the Calyx expressing Kr, detected by the rabbit Kr antibody. (D, E) Images showing the MB cell body region in larval brains expressing the Kr::GFP reporter. Kr::GFP is shown in green, DNA in grey, and Mira in red, with accompanying mono-coloured images. Asterisks mark MBNBs, while arrows indicate two cells adjacent to the Calyx expressing Kr::GFP. Kr::GFP signals were detectable in MBNBs and their adjunct cells, as well as cells attached to the Calyx. All scale bars: 20 µm.

**Table 5.** Adult central brains with EdU-positive clones overexpressing Kr in neuroblasts.

| Genotypes | EdU | Total | Ratio |
| --- | --- | --- | --- |
| *Elav, gal80ts > Kr::V5* | 0 | 14 | 0.0% |
| *Elav, gal80ts > GFP* | 0 | 14 | 0.0% |
| *wor, gal80ts > Kr::V5* | 0 | 15 | 0.0% |
| *wor, gal80ts > GFP* | 0 | 6 | 0.0% |
| *insc, gal80ts > Kr::V5* | 0 | 4 | 0.0% |
| *insc, gal80ts > GFP* | 0 | 6 | 0.0% |

Number of adult brains with EdU-positive clones and the total number of brains examined.

### Kr is expressed in MBNBs during postembryonic development

Our findings suggest a specific role for *Kr* in regulating the proliferative capacity and neurogenic activity of MBNBs. Kr is implicated in diverse developmental processes in *Drosophila*. Beyond its well-known role as a gap gene in early embryonic segmentation^52^ ^53^, Kr also serves as one of tTFs, controlling the developmental timing of neural fate decisions within the central nervous system^24,54,55^. However, its role in neurogenesis beyond the embryonic stage and in specific NB lineages, including MBNBs, remains poorly understood.

To explore the role of Kr in MBNBs, we examined its expression pattern specifically in MBNBs during various developmental stages. To detect the endogenous expression of Kr proteins, we obtained previously published two Kr-specific antibodies^56,57^. Using these antibodies, we were able to detect Kr expression in many NB lineages in wild type embryos as well as its ectopic expression in the *If* mutant eye imaginal discs (Fig. S6A, S7), as reported previously^24,27^.

We first examined Kr expression in MBNBs in developing embryos. It has been shown that the four MBNBs are individually specified in the procephalic neuroectoderm after embryonic stage 9^58^. In embryos in embryonic stages later than stage 12, we were able to clearly identify MBNBs based on the co-expression of Mira and mCD8::GFP induced by *OK107-*gal4 (Fig. S7). In these late embryos, we observed many cells expressing Kr scattered in their developing brains (Fig. S7). However, in both wild type and *If* mutants, Kr expression was not detected in MBNBs (Fig. S7A-C) and only occasionally observed in a non-MBNB cell in MBNB lineages per hemisphere (Fig. S7C).

We also examined Kr expression in postembryonic stages. Similar to embryos, we observed Kr expression in numerous neurons across the central brain in third instar larval brains (Fig. 5A, B). However, in the larval brains, weak Kr signals were detected in MBNBs, marked by Mira and mCD8::GFP induced by *OK107-*gal4 or *mb247*-gal4, using both Kr antibodies (Fig. 5A, B). Additionally, stronger Kr expression was observed in a few cells in MB lineages near the Calyx (Fig. 5C). To validate these findings, we also utilised a Kr::GFP reporter^59^. This reporter line carries a bacterial artificial chromosome containing an approximately 88-kb region on the second chromosome (2R: 25168930 to 25256753) encompassing the entire Kr gene locus (2R:25226611 to 25231394), in which the Kr gene is fused to a GFP-FLAG-PreScission-TEV-BLRP tag sequence at its 3’ end and thus is designed to reflect its native expression pattern^59^. Indeed, this reporter construct successfully rescued the viability of a Kr null mutant (*Kr^1^/Df(2R)Kr10*), confirming its functionality. In line with observations from Kr antibody staining, the Kr::GFP reporter exhibited weak GFP signals in MBNBs and more pronounced GFP signals in MB neurons adjacent to the Calyx in larval brains (Fig. 5D, E). The reporter also highlighted GFP signals in cells close to MBNBs, likely representing GMCs or immature MB neurons that might inherit Kr::GFP from MBNBs (Fig. 5E). These results indicate that Kr is expressed in MBNBs during larval stages. Supporting this, tissue-specific RNAseq data from FlyAtlas2 and modENCODE, as well as MBNB-specific RNAseq analysis, also revealed Kr expression in the larval central nervous system and a modest expression in larval MBNBs^60,61^.

We also examined Kr expression in pupal and adult brains. In pupal brains, the number of Kr-expressing cells were significantly reduced and specific Kr signals were undetectable in MB lineages (Fig.S8A). In adult brains, while no MBNBs were present, we occasionally observed cells expressing weak nuclear Kr signals in MB cell bodies (Fig. S8B). Additionally, there were a number of cells exhibiting cytoplasmic Kr signals co-localised with strong Mira signals above the MB cell body, whose identity remains unclear (Fig. S8C, D).

We further tested the impact of MBNB-specific depletion or overexpression of Kr on MBNBs using *OK107-*gal4 driver. The induction of *Kr::V5* led to high rate of lethality, similar to general NB drivers, yet allowing the production of a small number of surviving adult flies. While these adult flies exhibited a severe defect in the MB morphologies (Fig. S9), no mitotic MBNBs were observed in these adults. Meanwhile, the flies inducing *KrdsRNA* in MBNBs develop normally, and neither mitotic MBNBs nor any significant defect in MB morphologies were observed in the adult brains (n = 12, Fig. S9). This lack of phenotypes is likely due to the relatively low gene induction of the *OK107-*gal4 driver in MBNBs compared to MB neurons in postembryonic stages (Fig. 3E, S8A).

### Persistent Imp expression in persisting MBNBs of *insc>KrdsRNA* and *If* mutant adult brains

Finally, we asked how Kr mediates the timely elimination of MBNBs prior to the completion of development. MBNB elimination is controlled by various cell-intrinsic and extrinsic factors during postembryonic stages^18,19^. Notably, two RNA-binding proteins, IGF-II mRNA binding protein (Imp) and Syncrip (Syp), are expressed in MBNBs in opposing temporal gradients to regulate their elimination as well as the neural fates of their progeny^62^. Imp is expressed earlier than Syp to promote MBNB proliferation and early neural fate such as γ neuronal fate, and, as Imp expression descends, Syp expression gradually increases to promote the production of late MB neural types such as α’/β’ neurons and MBNB cell cycle exit and elimination^47,60^.

We analysed the expression patterns of Imp and Syp in *insc>KrdsRNA* and *If* mutant adult brains. Notably, there was a substantial expansion of the cell populations expressing Imp, where active MBNBs were embedded, in the dorsoposterior regions of *insc>KrdsRNA* adult brains as well as *If* mutant adult brains compared to control brains (Fig. 6A). While the size of Syp-expressing cell populations in these regions were not changed significantly, the expression levels of Syp in these domains appeared to be reduced in *insc>KrdsRNA* brains than in controls (Fig. 6B). We next assessed Imp and Syp expression specifically in persisting MBNBs in *insc>KrdsRNA* and *If* mutant adult brains. Intriguingly, almost all these MBNBs were found to continue expressing Imp (Fig. 6C, D, E), while co-expressing Syp (Fig. 6C, F, G). These observations suggest that Kr may regulate the timely elimination of MBNBs through its influence on Imp expression.

**Figure 6.**
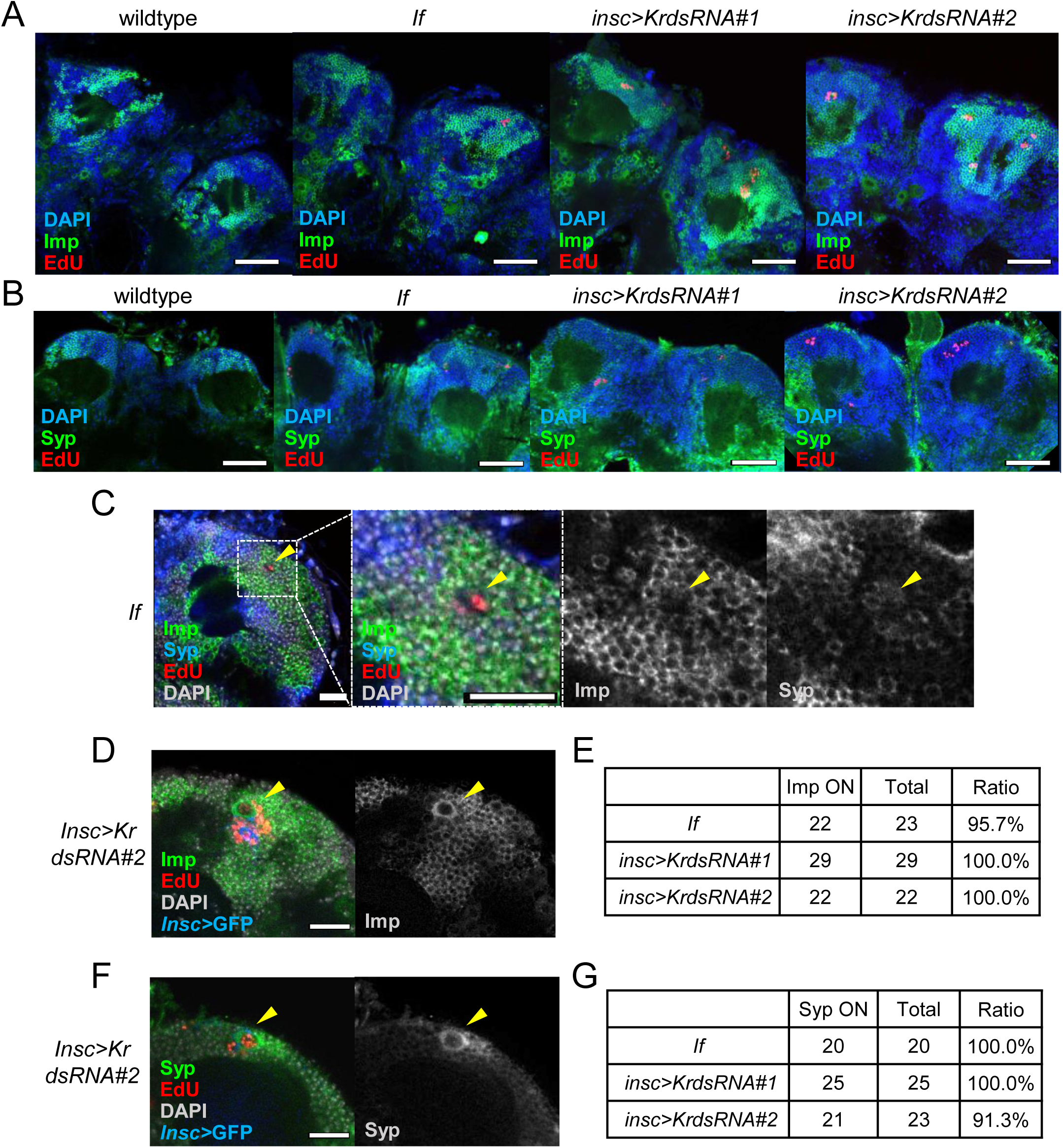
Continued Imp Expression in persisting MBNBs of *insc>KrdsRNA* and *If* mutant adult brains. (A, B) Representative images from the dorsoposterior regions of wild type, *If* mutant and *insc>KrdsRNA* adult brains, where MB cell bodies are located. Active MBNBs and their progeny were visualised by EdU labelling. Imp (A) or Syp (B) were detected using specific antibodies and shown in green, DAPI in blue, and EdU signals in red. Scale bars: 50 µm. Imp-expressing domains were significantly larger in *If* mutant and *insc>KrdsRNA* brains compared to control (A), while Syp-expressing regions remained largely unchanged (B). (C) Imp and Syp expression in active MBNBs in *If* mutant adult brains. In merged images, Imp is displayed in green, Syp in blue, EdU in red, and DAPI in grey, with accompanying mono-coloured views for each signal in grey. Arrowheads point to active MBNBs identified by EdU signals and cell size. (D, F) Imp and Syp expression in active MBNBs in *insc>KrdsRNA* adult brains. In merged images, Imp (D) or Syp (F) is shown in green, EdU in red, and DAPI in grey, *insc-*gal4-driven GFP signals in blue, with accompanying mono-coloured views for each signal in grey. Arrowheads highlight active MBNBs identified by EdU incorporation and GFP. Scale bars indicate 20 µm. (E, G) Number of MBNBs expressing Imp (E) and Syp (G) were quantified in *If* mutant and *insc>KrdsRNA* adult brains. Almost all the MBNBs found in these brains co-expressed Imp with Syp.

## Discussion

The adult brain of all animals is predominantly post-mitotic, with neurogenesis continuing in only a few specific regions, such as subgranular zone of the mammalian hippocampus^1,2^ and the optic lobes in *Drosophila*^15^. Our findings that neither mitotic cells nor EdU incorporation were detected and that nearly all cells displayed G1 cell cycle markers in control *Drosophila* adult central brains (Fig. 1, 4A) confirm this post-mitotic state. These observations are consistent with previous studies and indicate that neurogenesis in the central brain is tightly regulated to prevent unnecessary, potentially disruptive, proliferation.

Although it has recently been established that the *Drosophila* adult brain also possesses neurogenic capacity, discrepancies remain regarding the identity and proliferative state of adult neural progenitors. Several studies have reported potential NBs expressing common NB markers or G2-phase cell cycle markers in the adult central brain^7,15^. Consistent with these reports, we observed cells labelled by Mira or insc-gal4-induced GFP and those expressing G2-specific signals of the Fly-FUCCI reporters (Fig. 1, 2). However, these cells typically expressed only one marker, making their identities unclear. Another study suggests the presence of quiescent neural progenitors in the adult central brains that can be activated upon acute brain injury to express a common NB marker Dpn and generate neurons^17^. Further characterisation using a common set of NB markers is needed to resolve these discrepancies and understand the regulation of adult neural progenitors.

Our bioinformatics analysis of CCRs using the recent scRNAseq data provided insights into the mechanisms maintaining the postmitotic state of the adult brains (Fig. 1C-E, S1-3). A marked suppression of most positive CCRs, such as cyclins and Cdks, in neuronal and glial cell clusters, along with relatively high expression of negative CCRs, such as *Wee1*, *fzr*, and *rux*, suggests a coordinated repression of cell cycle reentry that likely prevents unintended proliferation and maintains neural integrity in the adult brain. Additionally, our analysis identified CCRs with expression patterns suggesting noncanonical roles in neurons and glia. The sustained high expression of E2F1 and its co-activator Dp, non-canonical CDKs, and components of the ubiquitin-proteasome pathway such as APC/C and SCF across NBs and neuronal clusters indicates their potential involvement in neural plasticity, transcription regulation, neuronal activity, and the maintenance of neural function. These findings underscore the utility of scRNAseq in uncovering cell type-specific functions of CCRs, providing deeper insights into the regulatory mechanisms preserving neural integrity and highlighting potential targets to explore adult neurogenesis and neural regeneration.

We observed the presence of proliferating MBNBs in the central brain of *If* mutant adult flies (Fig. 2, 3), marking the first example of a spontaneous mutation that modulates the neurogenic capacity of adult animals. These MBNBs continue proliferating over three weeks (Table 1, Fig. 4C). This prolonged activity suggests a fundamental change in their developmental trajectory, not merely a delay in their typical elimination. Notably, they maintain their stem cell properties, undergoing asymmetric cell divisions to produce differentiated daughter cells, as most clones contain single NBs (Fig. 2, 3). Given that MBNBs are the last to be eliminated during development^9,12^, it is highly plausible that in *If* mutants, these cells evade the usual elimination process at the late pupal stage and persist into adulthood. Nevertheless, our current data cannot rule out the possibility that these cells could arise from previously unidentified quiescent progenitors in the adult brain. Further research is necessary to definitively determine the origins and characteristics of these persisting MBNB-like cells in the *If* mutant adult brain.

Our findings reveal a novel role for a conserved transcription factor Kr in regulating the survival and proliferation of MBNBs in the *Drosophila* adult brain. Building on previous reports that linked the *If* mutation to the misexpression of *Kr*^27,28^, we demonstrate that dysregulation of *Kr* expression, whether through upregulation or downregulation, leads to abnormal MBNB activity. Specifically, we observed that the number of persistent MBNBs in *If* mutant adult brains was strongly suppressed by reducing Kr levels in NBs through *Kr*shRNA induction (Fig. S5). Conversely, depleting Kr proteins using *Kr*dsRNA led to the retention of active MBNBs in a wild-type background (Fig. 4). These findings highlight the importance of precise regulation of Kr expression levels in the timely cell cycle exit and elimination of MBNBs during development.

Our results suggest that the function and regulation of Kr in MBNB control is likely complex. The *If* mutation, a dominant-negative *Kr* allele where Kr has been shown to be ectopically expressed (Fig. S6A)^27,28^, leads to abnormal neuroblast activity in adult brains (Fig. 2, 3). Our data showing a rescue of this phenotype by Kr reduction through *Kr*shRNA (Fig. S5) suggest that *Kr* upregulation causes MBNB retention. However, simply overexpressing Kr in NBs using NB-specific drivers did not lead to MBNB persistance (Table 5). Moreover, our result with Kr depletion using *insc*>*Kr*dsRNA in the wild type flies suggests that the loss of Kr also induces MBNB retention (Fig. 4, S5). These results indicate a multifaceted role of Kr, involving both the promotion and inhibition of MBNB elimination under different contexts. Indeed, it has been shown that Kr has multiple roles in neuronal patterning, including in neurons^63^, consistent with our observation of its expression in MB neurons (Fig. 5, S7, S8).

We further investigated the mechanism by which Kr regulates MBNB elimination. Given the role of Imp in promoting NB survival and proliferation and Syp in promoting NB cell cycle exit and elimination^47,60^, we examined the impact of Kr on their expression levels. We found that Imp continues to be highly expressed in MB cell body regions in both *insc>KrdsRNA* and *If* mutant adult brains, compared to wild type (Fig. 6). The sustained MBNBs in these brains also exhibit continuous high expression of Imp (Fig. 6C-E), suggesting that Kr may regulate timley cell cycle exit and eliminianation of MBNBs through the regulation of Imp expression in MBNBs.

Interestingly, Syp is still expressed and often co-expressed with Imp, albeit at slightly reduced levels in the *insc>KrdsRNA* condition (Fig. 6B, C, F, G). This is somewhat unexpected given the previously reported antagonistic relationship between Imp and Syp^47,53,60^. However, this antagonism appears to be not strict: RNAi-mediated downregulation of one leads to upregulation of the other, but overexpression does not necessarily downregulate the other^47,60^, consistent with our observation of the co-expression of Imp and Syp in the persisting MBNBs (Fig. 6C-G). Persistent Syp expression may explain why MB structures appear unaffected in *If* mutants despite Imp upregulation (Fig. S4), as the Syp program may still be operating. While our data suggest that Kr regulates MBNB proliferation primarily through Imp, it is still possible that Kr may impact other factors that are known to regulate MBNB elimination, such as Chinmo, E93, and Rx^48,64,65^, contributing to the observed phenotype.

Kr is one of the tTFs in many NBs, regulating NB temporal patterning and proliferation. While Kr’s direct role in NB elimination has not been examined, late tTFs such as Cas and Svp, expressed during larval stages, directly regulate NB cell cycle exit and elimination^66^. Since tTFs cross-regulate each other^18,54^, Kr might impact MBNB elimination by modulating the expression or activity of later tTFs. However, the role of Kr in MBNB regulation is likely distinct from its function as a tTF. It was shown that Kr knockdown in embryonic NBs skips the Kr window without affecting the expression of later tTFs or the production of other neural types^24^. Additionally, our analysis of Kr expression in MBNBs showed that Kr is not strongly induced in MBNBs at any developmental stage and is expressed at a low level in MBNBs during larval stages (Fig. 5, S6, S7). Moreover, the impact of Kr manipulation is specific to MBNBs; while we induced Kr depletion using *insc>KrdsRNA*, which should impact all NBs, no obvious defects were observed in the elimination of other NBs.

The key question remains: How does Kr regulate Imp expression? Imp continues to be expressed in the persisting MBNBs (Fig. 6), indicating Kr’s cell-autonomous regulation. Our data and previous MBNB-specific RNAseq data^61^ show that Kr is expressed in MBNBs in larval brains, albeit weakly (Fig. 5). Considering its low expression, Kr may play a more permissive role in MBNBs, unlike the inductive role known for tTFs. As a transcription factor, Kr might directly regulate Imp transcription by binding to its promoter or influencing chromatin structure. Alternatively, Kr could indirectly regulate Imp expression by altering cell fate or acting in other NBs to influence MBNBs non-cell autonomously. MBNBs exhibit unique behaviours, such as producing only three types of neurons, not arresting during the embryo-to-larva transition and being the last NBs to be terminated^9^, suggesting unique regulatory mechanisms in which Kr may play a part. Further investigation is needed to elucidate the precise role and regulation of Kr in MBNBs.

Our results, together with mounting evidence, point to the high plasticity and adaptability of the neurogenic capacity of the adult brain. Intriguingly, it is known that some insects, such as certain bees and ants, continue to exhibit MBNB proliferation in adult brains^67,68^, similar to what we observed in the *If* mutant and *insc>KrdsRNA* fliles. Sustained neurogenesis and regenerative capacity in the adult brain present clear advantages. However, such proliferative potential must be balanced with the risk of interfering with the existing neural network and spontaneous tumorigenesis. Therefore, the mitotic and regenerative activity and the population size of neural progenitors in the adult brain must be tightly regulated. While we observed that newly generated neurons are integrated into the existing MB structures (Fig. 3), it remains unclear whether these newly generated neurons are functional and can contribute to MB function. *If* mutants and *insc>KrdsRNA* flies develop without significant delay and are mobile and fertile. The overall structures of the MBs and the adult brain appear to be unaffected in the *If* mutant (Fig. S4). However, there may be subtle alterations in the detailed structure and/or function of the MBs in these flies. It may be of interest to carefully examine the morphology, behaviour, and learning ability of these flies.

Notably, one of the Kr orthologues in mammals, KLF4, is a member of the pluripotency factors^69^. Cyclic expression of these factors, including KLF4, in mouse NSCs inhibits aging phenotypes in long-term in vitro culture^70^. Another Kr orthologue, KLF9, is expressed in NSCs in the hippocampus of adult mice to maintain NSC quiescence, and its forced expression in mature dentate granule cells promotes the activation of adult NSCs in a non-cell-autonomous manner^71,72^. Thus, Kr family proteins play a crucial role in controlling the longevity and proliferative capacity of NSCs across invertebrate and vertebrate species. Further studies on the role of Kr family transcription factors using various models will provide deeper insights into the mechanisms that govern the regenerative ability of the adult brain and may offer a window of opportunity to enhance neurogenic and regenerative capacity through their genetic or pharmacological manipulation.

### Limitation of the study

Our study has several limitations. First, our cell cycle analysis and scRNAseq data, while informative, may lack the sensitivity to detect minor populations of slow-dividing cells or active neural progenitors. Second, the precise impact of the *If* mutation on Kr expression remains unknown, and our study did not fully elucidate its effects on Kr transcription or the adult brain structure and function. Our data with *If* mutation and *insc>KrdsRNA* suggest a complex role for Kr, as both its upregulation and downregulation result in abnormal MBNB activity. While our data imply a cell-autonomous effect, *Kr* misregulation in other NBs might also impact MBNBs non-cell autonomously. Third, the temporal dynamics of *Kr* expression and environmental influences, such as temperature and nutrition, were not fully explored and may affect Kr’s regulatory role in MBNBs. Fourth, while we identified a significant role for Imp in Kr-mediated MBNB regulation, the exact mechanisms remain unclear. Kr might regulate Imp directly or indirectly, and other factors could also be involved. Finally, our findings in *Drosophila* may not directly translate to other species. Genetic and epigenetic contexts could influence Kr function differently across organisms, necessitating further comparative studies.

## Author contributions

Conceptualisation, Y.K.; Methodology, J.M., X.S., and Y.K. ; Software, H.S.; Investigation, J.M., X.S., X.X., H.S., and Y.K.; Resources, J.M., X.S., Y.A., and Y.K; Writing – Original Draft, X.S., H.S., and Y.K., ; Wirting – Review & Editing, J.M., Y.A., and Y.K.; Visualisation, J.M., X.S., H.S., and Y.K. ; Supervision, Y.K. ; Funding Acquistion, Y.A. and Y.K.

## Acknowledgements

We thank Dr Margaret HO Ho for discussions, for sharing *Drosophila* reagents and for providing the initial training for the dissection and immunostaining of *Drosophila* adult brains, Dr Wei Wu at Shanghai Fly Center, Dr Simon Collier at Cambridge *Drosophila* Facility at ShanghaiTech University for the help with maintenance and transporting *Drosophila* stocks and for embryo injection services, Dr. Reinhard Klug and the VDRC for the assistance with the transportation of *Drosophila* stocks and RNAi fly lines, and Dr Cédric Maurange, Dr Nan Liu and Dr Yukinori Hirano for discussions, and all *Drosophila* colleagues at ShanghaiTech University, in particular, Dr Ji-long Liu, Dr Jingnan Liu and Dr Guanjun Guo, and China *Drosophila* community for discussion, sharing information and reagents. We thank Ilan Davis, Chris Rushlow, Claude Desplan, Kuniaki Saitoh, Rita Sousa-Nunes and Kei Ito for antibodies and technical advice. We also thank all Kimata lab members at ShanghaiTech University and the University of Cambridge for cooperation and discussions. This work was supported by the ShanghaiTech University start-up grant (2018F0202-000-06), National Natural Science Foundation of China (NSFC) Mainshang grant (32170746) and Research Fund for International Scientists (RFIS, 32150710520) and to YK.

## Materials and Methods

### Fly stocks and culture

The *Drosophila* lines used in this study are as follows. Gal4 driver lines specific for the mushroom body neurons, *mb247-gal4* (a gift from Dr Margaret Ho) and *OK107-gal4; UAS-mCD8::GFP* (a gift from Dr Yi Zhong). Gal4 driver lines for NBs, *insc*-*gal4* (a gift from D Andrea Brand), *UAS-dicer2*; *insc*-*gal4, UAS-mCD8::GFP*/CyO (a gift from Dr Eugen Knoblich), *wor-gal4* (a gift from Dr Cahir O’Kane). An eye-specific Gal4 line *ey-gal4*^73^. *Kr*dsRNA lines (v104150, v40871) and an *E2F2*RNAi line (v100990) from VDRC. A *Kr*shRNA line (THU3653) and an *Hb*shRNA line (THU5932) from TsingHua Fly Center. *pTub-gal80ts* lines (BL7107, BL7108), a GFP reporter line, *UAS-mCD8::GFP* (BL5130), a Kr overexpression line *UAS-Kr-V5* (BL83301^50^), a loss-of-function mutant and a deficiency line, *Kr*^1^ (BL3494) and *Df(2R)Kr10* (BL4961), and a Kr protein reporter line *PBac{Kr-GFP.FPTB}* (BL56152) from the Bloomington Drosophila Stock Center. FlpOUT-Gal4 lines, *hsflp[122]; pAct>CD2>gal4; UAS-dicer2, UAS-GFP* and *hsflp, pAct>CD2>gal4, UAS-RFP* (gifts from Dr Ji-long Liu and Dr Chenhui Wang, respectively). A Fly-FUCCI line *pUbq-GFP-E2F1, mRFP-NLS-CycB* (a kind gift from Dr Bruce Edgar). A double balancer line, *If/Cyo; Sb/TM6B,* and wild type and control stocks, *Oregon R* and *w^1118^* (Cambridge Drosophila Facility). An additional *If* mutant stock *w^1118^; If/CyO* (BCF360 from Shanghai Fly Center).

All *Drosophila* lines were maintained and cultured with common cornmeal agar media, with 12 hr-12 hr day-night cycles, and at 50-70% humidity. All fly experiments were conducted at 25 ℃ except for RNAi and over-expression experiments in which flies were grown at 29°C for higher Gal4-dependent induction, as specifically noted. 20 to 30 flies were contained in each tube and flipped them into new tubes every two to three days. Fly crossing was done with around 10 virgins and 8 males with specific genotypes in each tube. In most crosses, virgins of Gal4 driver lines were collected and crossed with the male flies with UAS elements. The progeny with the specific genotypes were selected based on visible genetic markers for analysis.

### Immunofluorescence and confocal imaging of *Drosophila* brains

*Drosophila* adult brains were dissected in 1xPBS following the protocol described in Wu and Luo, 2006^74^. The brains were then fixed in freshly made fixative containing 8% formaldehyde, 1xPBS, 0.5mM EGTA and 5mM MgCl_2_, at room temperature for 1 hour. The fixed brain samples were then washed in PBST (1xPBS, 0.3% Triton-X), blocked with 3% BSA in PBST (PBSTB) at room temperature for 1 hour, and then incubated in PBSTB containing appropriate primary antibodies at 4°C overnight. After washes, the samples were further incubated in PBSTB containing secondary antibodies and DAPI (Cell Signaling Technology) at room temperature for 3 hours, then washed in PBST and kept in mounting media (Vectashield, VectorLab). The brain samples were then mounted onto microscope slides and kept in 4℃ till imaging. As for pupal brains, the whole body of pupae with its shell removed were fixed in freshly made fixative containing 8% formaldehyde, at room temperature for 40 minutes, and after washing fixed pupal brains were then carefully dissected in 1x PBS. In larvae, eye-brain complexes were fixed in freshly made fixative containing 4% formaldehyde, 1xPBS, 0.5mM EGTA and 5mM MgCl_2_, at room temperature for 30 minutes. After washing, blocking, and staining steps mentioned as above, larval brains were been carefully separated from the complexes and mounted onto slides.

For embryos, parental flies were incubated in a small cage with an apple juice agar plate pasted with full of yeast extract paste, and replaced the old plate with a new plate with yeast every day. Collecting eggs at the desired stage of development by brushes and transfer them to a small basket. Dechorionate eggs by putting them in 50% bleach and wash the dechorionated eggs by water. The dechorionated embryos were then transfered into a glass bottle (1ml/6ml) containing 8%Flormaldehyde/PBS: Heptane (1:1) and incubated at room temperature for 20min. Incubating embryos 1x volume of −20℃ MeOH for 1min after removing the lower FA phase. Washed the embryos with 1ml MeOH for 5min 3 times. The embryos were then rehydrated by 15min PBS and PBST (PBS +0.05%TritonX-100) incubation respectively. Immunostaining steps were the same as mentioned above, except embryos were blocked in PBTA(PBS +0.05%Triton X-100 + 1%BSA) at room temperature for 1 hr.

Images were taken by a Nikon C2 confocal microscope in four channels with emission laser wavelengths, 405, 488, 568 and 647 nm. Image reconstruction and analysis were processed and conducted using NIS-Elements AR 5.20 software and Image J ImageJ^75^. All projected images shown in this paper were processed by maximum intensity projection. For better visualisation, a 0.5 Gaussian blur filter was applied to every image shown.

### Antibodies

Primary antibodies used in this study are as follows: rabbit Phospho-Histone H3 (Ser10) antibody (Invitrogen PA5-17869, 1:400), rat anti-Elav (Developmental Studies Hybridoma Bank, 7E8A10 1:50,), rat anti-Mira (1:400) and rat anti-Dpn (1:100, kind gifts from Dr Chris Doe), rabbit anti-Kr antibody (a gift from Dr Chris Rushlow, 1:1000)^56^, rat anti-Kr antibody (from Asian Distribution Center for Segmentation Antibodies, 1:100)^57^, guinea pig anti-Syp (a gift from Ilan Davis, 1:200), rat anti-Imp (a kind gift from Dr Claude Desplan, 1:200), mouse anti-FasII (DSHB, 1D4, 1:40), rabbit anti-Cleaved *Drosophila* DCP-1 (Cell Signaling Technology #9578, 1:100), mouse anti-V5 antibody (Invitrogen, SV5-Pk1). Second antibodies used in this study are goat anti-mouse, rat or rabbit secondary antibodies conjugated with Alexa Flour 488, 568 or 647 (Invitrogen, 1:500), DAPI (Cell Signaling Technology, 1:1000).

### EdU labelling of proliferating cells in *Drosophila* adult brains

EdU labelling was performed to detect proliferating cells in *Drosophila* adult brains, following the protocol outlined by Siegrist et al. (2010)^12^. EdU powder was dissolved in a 5% sucrose solution to achieve a final concentration of 0.5 mg/ml, and this solution was applied to paper to replace the regular food source in the culture vials. One-to three-day-old young adult flies were then collected and raised until the flies reached certain ages, and then fed with EdU-supplemented food for a period of three days prior to brain dissection. For EdU labelling chase experiments, EdU exposure was ceased after the initial labelling period, and the flies were subsequently maintained for an additional three, seven, or fourteen days before brain dissection.

Following dissection, adult brains were subjected to immunofluorescence staining using standard procedures, with the exception of an additional step for EdU detection. EdU Click labelling was performed in the dark for 30 minutes prior to primary antibody incubation, following the manufacturer’s instructions (APExBIO EdU Imaging Kits Cy3, #K1075).

EdU-incorporated cell clones were visualised and manually counted in adult brains using a Nikon Ti2 fluorescence microscope equipped with a 20x objective lens. Statistical analyses were conducted using GraphPad Prism software. Data distribution was first assessed with the D’Agostino-Pearson omnibus normality test. For data that followed a normal distribution, an unpaired t-test was employed. For non-normally distributed data, the Mann-Whitney U test was utilized. Differences were considered statistically significant at p < 0.05. Significance levels were denoted as follows: *p ≤ 0.05, **p ≤ 0.01, ***p ≤ 0.001, ****p < 0.0001; ns indicates non-significant results.

### Differential Expression Analysis of Cell Cycle Genes in the *Drosophila* Adult and Larval Brains

The original data files of the single-cell RNA sequencing of *Drosophila* adult brains^32^ were obtained from the NCBI Gene Expression Omnibus (series number GSE107451) and processed using Seurat v3^76^. To ensure data quality and remove potential dying cells, we restricted the dataset to samples with 200 to 3700 unique genes and a UMI-to-gene ratio smaller than 6. The resulting dataset comprised 56,619 samples and 12,925 features. For normalisation, raw counts were first adjusted to account for sequencing depth by scaling each cell by the total number of counts and multiplying by a scaling factor of 10,000. The normalised counts were then log-transformed using the natural logarithm (log1p), which transforms the data using the formula log(1 + x), where x represents the normalized counts. Principal Component Analysis (PCA) was conducted on the 2000 highly variable genes identified. Shared Nearest Neighbour (SNN) cell clustering at a resolution of 0.2 or 8 was performed using the first 20 principal components, and t-SNE reduction was applied to visualise the clustering results. Based on the differential expression of the cluster markers and comparison with the original annotations^32^, 17 or 84 cell clusters were identified, respectively. The expression of 112 CCR genes across these clusters was then visualised using dot plots. In these plots, the colour intensity of each dot represents the average expression level of the CCR gene, with darker red indicating higher expression levels. The size of each dot indicates the proportion of cells within the cluster expressing the gene, with larger dots representing higher proportions. To illustrate the heterogeneity within clusters, heatmap plots were generated for each CCR gene. In these heatmaps, each gene is represented by a row and each cell by a short line, highlighting the variability of expression levels among individual cells within clusters.

For the analysis of *Drosophila* larval brains, the original data files of the single-cell RNA sequencing of larval brains^33^ were obtained, processed and presented similar to the adult brain. The analysis dataset contains 4349 samples and 12,942 features. A resolution of 2 was used and 29 clusters were identified.

### The rescue of the adult neurogenic phenotype of the *If* mutant by *Kr* RNAi

To test if NB-specific knockdown of Kr or Hb can rescue the ectopic NBs in the *If* mutant adult brain, the transgenic lines of *UAS-KrshRNA/TM3, Sb* and homozygous *UAS-HbshRNA* were first crossed with the double balancer stock, *w^1118^; If/CyO; Sb/TM6B, Tb*, and the male progeny with the genotypes of *w^1118^; If/+; UAS-KrRNAi/TM6B, Tb* and *w^1118^; If/+; UAS-HbshRNA/TM6B, Tb* were obtained. Next, these males, together with the males of control *w^1118^* and *w^1118^; If/CyO* were crossed with *w^1118^, UAS-dicer2; insc-gal4, UAS-mCD8::GFP/CyO* virgin females, and all progeny from these crosses were raised at 29°C throughout. 3-day-old adult flies with the following genotypes: 1) *w^1118^, UAS-dicer2; insc-gal4, UAS-mCD8::GFP/+*, 2) *w^1118^, UAS-dicer2; insc-gal4, UAS-mCD8::GFP/If*, 3) *w^1118^, UAS-dicer2; insc-gal4, UAS-mCD8::GFP/If*, *UAS-KrshRNA/+*, 4) *w^1118^, UAS-dicer2; insc-gal4, UAS-mCD8::GFP/If*, *UAS-HrshRNA/+*, were collected and dissected to analyse adult brains by immunostaining.

Additionally, the transgenic lines of *UAS-KrdsRNA/CyO* were first crossed with *w^1118^; If/CyO,* and then the female progeny with the genotypes of *w^1118^*; *If/UAS-KrdsRNA* and *w^1118^; If/UAS-Kr-V5* were obtained. Next, these females, were first crossed with *w^1118^; If/CyO* again. Finally, male progeny with the genotypes of *w^1118^; If, UAS-KrdsRNA/CyO* and *w^1118^*; *If, UAS-Kr-V5/CyO,* together with *w^1118^* and *w^1118^; If/CyO,* were crossed with *w^1118^, UAS-dicer2; insc-gal4, UAS-mCD8::GFP/CyO* females, and all progeny from these crosses were raised at 29°C throughout. 3-day-old adult flies with the following genotypes: 1) *w^1118^, UAS-dicer2; insc-gal4, UAS-mCD8::GFP/+,* 2) *w^1118^*, *UAS-dicer2; insc-gal4, mCD8::GFP/If*, 3) *w^1118^, UAS-dicer2; insc-gal4, UAS-mCD8::GFP/If, UAS-KrdsRNA,* were collected and dissected to analyse adult brains by immunostaining.

### Kr over-expression in NBs

To overexpress Kr in NBs, the crossing between *UAS-Kr-V5* line and either *insc-gal4*, *wor*-*gal4*, and *Elav-gal4* did not yield any third instar larva while hatched eggs were observed, indicating that continuous Kr overexpression in NBs causes lethality in early larval stages. We used *insc-gal4*, *wor*-*gal4*, and *Elav-gal4* ga80ts and crossed with *UAS-Kr-V5* to allow survival and development into adult flies. Progeny was heatshock 3 days after crossing to induce UAS-Kr-V5, adult brains were fed with EdU and the adult brains were dissestede and immunostained for analysis. No EdU incorporation in any of the adult brains n>20.

## Supplementary information

**Figure S1 Heatmap of CCR gene expression in 17 clusters of *Drosophila* adult brain**

The heatmap displays the expression levels of 112 selected CCR genes across 17 identified clusters in the *Drosophila* adult brain. Each row represents a CCR gene, and each column corresponds to an individual cell within the clusters. The colour intensity indicates the expression level, with darker red indicating higher expression levels and lighter shades indicating lower expression levels. This visualization highlights the heterogeneity of CCR gene expression within each cluster.

**Figure S2 Dot plot and heatmap of CCR gene expression across clusters of the *Drosophila* larval brain.**

(A) Dot plot showing the expression patterns of CCR genes across 29 cell clusters identified through Seurat using the recent single-cell RNA sequencing data from *Drosophila* larval brains^33^. The colour intensity of each dot represents the average expression level of the CCR gene, with darker red indicating higher expression levels similar to Fig. 1D. The size of each dot indicates the proportion of cells within the cluster expressing the gene, with larger dots representing higher proportions.) and dot size representing the proportion of cells expressing each gene. (B) Heatmap displaying the expression levels of CCR genes in *Drosophila* larval brains, presented similarly to Fig. S1. Each row represents a CCR gene, and each column corresponds to an individual cell, with colour intensity indicating the expression level.

**Figure S3 Dot plot of CCR gene expression across the 84 clusters of the *Drosophila* adult brain, identified in Davie et al., 2018**^32^.

The dot plot illustrates the expression of 112 selected CCR genes across 84 identified clusters in the *Drosophila* adult brain. Each dot represents the average expression level of a CCR gene within a cluster, with the colour intensity indicating the expression level (darker red indicates higher expression). The size of each dot represents the proportion of cells within the cluster expressing the gene, with larger dots indicating higher proportions. This plot provides an overview of the distribution and prevalence of CCR gene expression across all identified clusters.

**Figure S4 The overall structures of the adult mushroom bodies are not altered in *If* mutants**

(A) The structures of the MBs were visualised by *mb247>mCD8::GFP* in *If* mutant (*If/CyO,* top) and control (*Sp/CyO*, bottom) adult brains. The anterior and posterior views of the adult brains, where MB lobes, calyx, and cell bodies are visible, are shown on the left and right, respectively. DAPI (blue) and GFP (green). Insets show magnified images. Scale bars: 50 µm. (B) Anterior views of the 3D-reconstructed confocal images of the central brains in wild type (top) and the *If* mutant (*If/CyO,* bottom) where MB lobes were visualised using anti-FasII antibody (green). No obvious morphological alterations were observed. Scale bars: 50 µm.

**Figure S5 Kr knockdown using *Kr*shRNA induction suppresses ectopic NB formation in *If* mutant adult brains.**

(A) Representative images of adult central brains from control *(+/+, insc>lacZ*), *If* mutant (*If/+, insc>lacZ*), *If* mutant with NB-specific *KrshRNA* induction (*If/+, insc>KrshRNA*), and *If* mutant with NB-specific *HbshRNA* induction (*If/+, insc-gal4>HbshRNA*). DNA is shown in blue, *insc*-gal4-driven mCD8::GFP in green, and pH3 in red. The numbers and ratios of brains exhibiting pH3-positive cells among the total number of brains examined are quantified and shown at the top. Scale bars: 50 µm. Mitotic NB formation in *If* mutant adult brains was partially suppressed by NB-specific induction of *KrshRNA*, but not *HbshRNA*. (B) EdU labelling was conducted on adult flies of the indicated strains for three to five days post-eclosion. Brains were subsequently dissected, and the number of EdU-positive cell clones per hemisphere was quantified and displayed in scattered dot plots. Means and standard deviations are represented by thick and thin red bars, respectively, with actual values displayed above the bars in red. ‘n’ denotes the number of brains analysed per condition. Statistical significance was determined using the Mann-Whitney U test, with significance levels denoted as: * (p ≤ 0.05), ** (p ≤ 0.01), *** (p ≤ 0.001), **** (p < 0.0001). ‘ns’ indicates non-significant results.

**Figure S6 The *KrdsRNA* constructs repress Kr expression more efficiently than the *Kr*shRNA construct**

(A) Eye imaginal discs from of wild-type (*+/+*) and *If* mutant (*If/+*) third instar larvae stained by DAPI (blue), anti-Kr antibodies raised in a rat (Rt, green) and a rabbit (Rb, red)^56,57^. Overexpression of Kr proteins was detected by both anti-Kr antibodies in differentiating cells in *If* mutant eye discs, but not in wild-type discs. Scale bars: 100 µm. (B) Eye imaginal discs of *If* mutants, either uninduced (*-*) or expressing *KrshRNA*, *KrdsRNA#1*, or *KrdsRNA#2* driven by the eye region-specific *eyeless*-gal4 (*ey-*gal4). Kr protein levels were detected using the rabbit anti-Kr antibody, verified for specificity and linearity previously^56^. DNA is shown in blue, and Kr signals in red. Scale bars: 100 µm. (C) Representative images of adult eyes of *If* mutants with no UAS construct, or *KrshRNA*, *KrdsRNA#1*, or *KrdsRNA#2* induced by *ey-*gal4. The *Kr*dsRNA constructs more effectively suppressed ectopic Kr expression in the eye imaginal discs of If mutants and more efficiently ameliorated morphological defects in the adult compound eyes than the *Kr*shRNA construct.

**Figure S7 Kr is undetectable in MBNBs in embryos**

(A, B) Representative images of the CNS in wild-type Stage 12 embryos. Kr proteins were detected using the rabbit anti-Kr antibody^56^. In merged images, DNA is shown in blue, *OK107-*gal4-driven mCD8::GFP in green, Kr in red, and Mira in grey, with accompanying mono-coloured images for each signal in grey. Scale bars: 20 µm. Asterisks indicate MBNBs, marked by the co-expression of *OK107>*GFP and Mira. (B) Arrows point to a neuronal cell in the MBNB lineage that shows clear Kr signals. While Kr is expressed in various cells within the embryonic CNS, it appears absent in MBNBs.

**Figure S8 Kr is undetectable in MBNBs in pupal and adult brains**

(A) Representative images of the dorsoposterior MB cell body region in pupal brains. In merged images, Kr-specific signals, detected using a rabbit anti-Kr antibody, are shown in red, DNA in grey, *OK107*>GFP in green, and Mira in cyan. Mono-coloured views are provided for Kr and Mira. Asterisks indicate MBNBs where Kr signals were not detectable. (B-D) Images of the dorsoposterior region of the wild-type adult brain, focusing on MB cell bodies and the Calyx. In merged images, Kr signals are shown in red, OK107-driven GFP in green, and Mira in cyan, with single-coloured views for each signal provided alongside the merged images. No MBNBs were detected in wild-type adult brains. (B) Shows a MB neuron with weak nuclear Kr signals (arrow). (C, D) Display uncharacterised cells near the MB cell body, exhibiting cytoplasmic Kr signals co-localised with Mira (arrows). Scale bars: 20 µm.

**Figure S9 Effects of Kr depletion or overexpression in MB lineages using *OK107*-gal4**

Posterior views of the 3D-reconstructed confocal images of the central brains of the control (no UAS gene), *OK107>KrdsRNA#2,* and *OK107>Kr::V5* adult flies where MBs were visualised by mCD8::GFP induced by *OK107*-gal4 (green). While the control and *OK107>KrdsRNA#2* adult brains did not exhibit any defects, Kr::V5 overexpression driven by *OK107-*gal4 caused high lethality rate and surviving adult flies displayed severe morphological defects in the MB structure. Scale bars: 50 µm.

## References

1. Negredo, P.N., Yeo, R.W., and Brunet, A. (2020). Aging and Rejuvenation of Neural Stem Cells and Their Niches. Cell Stem Cell 27, 202–223. 10.1016/j.stem.2020.07.002.

2. Niklison-Chirou, M.V., Agostini, M., Amelio, I., and Melino, G. (2020). Regulation of Adult Neurogenesis in Mammalian Brain. Int J Mol Sci 21. ARTN 4869 10.3390/ijms21144869.

3. Doe, C.Q. (2008). Neural stem cells: balancing self-renewal with differentiation. Development 135, 1575–1587. 10.1242/dev.014977.

4. Casas Gimeno, G., and Paridaen, J. (2022). The Symmetry of Neural Stem Cell and Progenitor Divisions in the Vertebrate Brain. Front Cell Dev Biol 10, 885269. 10.3389/fcell.2022.885269.

5. Morales, A.V., and Mira, H. (2019). Adult Neural Stem Cells: Born to Last. Front Cell Dev Biol 7, 96. 10.3389/fcell.2019.00096.

6. Urbán, N., Blomfield, I.M., and Guillemot, F. (2019). Quiescence of Adult Mammalian Neural Stem Cells: A Highly Regulated Rest. Neuron 104, 834–848. 10.1016/j.neuron.2019.09.026.

7. Li, G.Y., and Hidalgo, A. (2020). Adult Neurogenesis in the Brain: The Evidence and the Void. Int J Mol Sci 21. ARTN 665310.3390/ijms21186653.

8. Truman, J.W., and Bate, M. (1988). Spatial and temporal patterns of neurogenesis in the central nervous system of Drosophila melanogaster. Dev Biol 125, 145–157. 10.1016/0012-1606(88)90067-X.

9. Ito, K., and Hotta, Y. (1992). Proliferation pattern of postembryonic neuroblasts in the brain of Drosophila melanogaster. Dev Biol 149, 134–148. 10.1016/0012-1606(92)90270-q.

10. Sousa-Nunes, R., Cheng, L.Y., and Gould, A.P. (2010). Regulating neural proliferation in the Drosophila CNS. Curr Opin Neurobiol 20, 50–57. 10.1016/j.conb.2009.12.005.

11. Maurange, C. (2012). Temporal specification of neural stem cells: insights from Drosophila neuroblasts. Curr Top Dev Biol 98, 199–228. 10.1016/b978-0-12-386499-4.00008-2.

12. Siegrist, S.E., Hague, N.S., Chen, C.H., Hay, B.A., and Hariharan, I.K. (2010). Inactivation of Both and Promotes Long-Term Adult Neurogenesis in Drosophila. Curr Biol 20, 643–648. 10.1016/j.cub.2010.01.060.

13. Yasugi, T., and Nishimura, T. (2016). Temporal regulation of the generation of neuronal diversity in Drosophila. Development, Growth & Differentiation 58, 73–87. 10.1111/dgd.12245.

14. von Trotha, J.W., Egger, B., and Brand, A.H. (2009). Cell proliferation in the Drosophila adult brain revealed by clonal analysis and bromodeoxyuridine labelling. Neural Dev 4, 9. 10.1186/1749-8104-4-9.

15. Fernández-Hernández, I., Rhiner, C., and Moreno, E. (2013). Adult Neurogenesis in Drosophila. Cell Rep 3, 1857–1865. 10.1016/j.celrep.2013.05.034.

16. Foo, L.C., Song, S., and Cohen, S.M. (2017). miR-31 mutants reveal continuous glial homeostasis in the adult Drosophila brain. The EMBO Journal 36, 1215–1226-1226. 10.15252/embj.201695861.

17. Crocker, K.L., Marischuk, K., Rimkus, S.A., Zhou, H., Yin, J.C.P., and Boekhoff-Falk, G. (2021). Neurogenesis in the adult brain. Genetics 219. ARTN iyab092 10.1093/genetics/iyab092.

18. Li, X., Chen, Z., and Desplan, C. (2013). Chapter Three - Temporal Patterning of Neural Progenitors in Drosophila. In Current Topics in Developmental Biology, A.E. Rougvie, and M.B. O’Connor, eds. (Academic Press), pp. 69–96. 10.1016/B978-0-12-396968-2.00003-8.

19. Maurange, C. (2020). Temporal patterning in neural progenitors: from Drosophila development to childhood cancers. Dis Model Mech 13. 10.1242/dmm.044883.

20. Chen, Y.-C., and Konstantinides, N. (2022). Integration of Spatial and Temporal Patterning in the Invertebrate and Vertebrate Nervous System. Front Neurosci-Switz 16. 10.3389/fnins.2022.854422.

21. Pearson, R., Fleetwood, J., Eaton, S., Crossley, M., and Bao, S. (2008). Krüppel-like transcription factors: a functional family. Int J Biochem Cell Biol 40, 1996–2001. 10.1016/j.biocel.2007.07.018.

22. Yuce, K., and Ozkan, A.I. (2024). The kruppel-like factor (KLF) family, diseases, and physiological events. Gene 895, 148027. 10.1016/j.gene.2023.148027.

23. Wieschaus, E., Nusslein-Volhard, C., and Kluding, H. (1984). Krüppel, a gene whose activity is required early in the zygotic genome for normal embryonic segmentation. Dev Biol 104, 172–186. 10.1016/0012-1606(84)90046-0.

24. Isshiki, T., Pearson, B., Holbrook, S., and Doe, C.Q. (2001). Drosophila neuroblasts sequentially express transcription factors which specify the temporal identity of their neuronal progeny. Cell 106, 511–521. Doi 10.1016/S0092-8674(01)00465-2.

25. Romani, S., Jimenez, F., Hoch, M., Patel, N.H., Taubert, H., and Jäckle, H. (1996). Krüppel, a Drosophila segmentation gene, participates in the specification of neurons and glial cells. Mech Dev 60, 95–107. 10.1016/s0925-4773(96)00603-x.

26. Pinto-Teixeira, F., and Desplan, C. (2016). Re-utilization of a transcription factor. Elife 5, e21522. 10.7554/eLife.21522.

27. Baker, N.E., Moses, K., Nakahara, D., Ellis, M.C., Carthew, R.W., and Rubin, G.M. (1992). Mutations on the second chromosome affecting the Drosophila eye. J Neurogenet 8, 85–100. 10.3109/01677069209084154.

28. Carrera, P., Abrell, S., Kerber, B., Walldorf, U., Preiss, A., Hoch, M., and Jäckle, H. (1998). A modifier screen in the eye reveals control genes for activity in the embryo. P Natl Acad Sci USA 95, 10779–10784. DOI 10.1073/pnas.95.18.10779.

29. Li, G.Y., Forero, M.G., Wentzell, J.S., Durmus, I., Wolf, R., Anthoney, N.C., Parker, M., Jiang, R.Y., Hasenauer, J., Strausfeld, N.J., et al. (2020). A Toll-receptor map underlies structural brain plasticity. Elife 9. ARTN e5274310.7554/eLife.52743.

30. Zielke, N., Korzelius, J., van Straaten, M., Bender, K., Schuhknecht, G.F.P., Dutta, D., Xiang, J.Y., and Edgar, B.A. (2014). Fly-FUCCI: A Versatile Tool for Studying Cell Proliferation in Complex Tissues. Cell Rep 7, 588–598. 10.1016/j.celrep.2014.03.020.

31. Technau, G.M. (2007). Fiber number in the mushroom bodies of adult Drosophila melanogaster depends on age, sex and experience. J Neurogenet 21, 183–196. 10.1080/01677060701695359.

32. Davie, K., Janssens, J., Koldere, D., De Waegeneer, M., Pech, U., Kreft, L., Aibar, S., Makhzami, S., Christiaens, V., González-Blas, C.B., et al. (2018). A Single-Cell Transcriptome Atlas of the Aging Drosophila Brain. Cell 174, 982-+. ARTN 998.e2010.1016/j.cell.2018.05.057.

33. Brunet Avalos, C., Maier, G.L., Bruggmann, R., and Sprecher, S.G. (2019). Single cell transcriptome atlas of the Drosophila larval brain. Elife 8. 10.7554/eLife.50354.

34. Park, D.S., Morris, E.J., Bremner, R., Keramaris, E., Padmanabhan, J., Rosenbaum, M., Shelanski, M.L., Geller, H.M., and Greene, L.A. (2000). Involvement of retinoblastoma family members and E2F/DP complexes in the death of neurons evoked by DNA damage. J Neurosci 20, 3104–3114. Doi 10.1523/Jneurosci.20-09-03104.2000.

35. Andrusiak, M.G., McClellan, K.A., Dugal-Tessier, D., Julian, L.M., Rodrigues, S.P., Park, D.S., Kennedy, T.E., and Slack, R.S. (2011). Rb/E2F regulates expression of neogenin during neuronal migration. Mol Cell Biol 31, 238–247. 10.1128/mcb.00378-10.

36. Castillo, D.S., Campalans, A., Belluscio, L.M., Carcagno, A.L., Radicella, J.P., Cánepa, E.T., and Pregi, N. (2015). E2F1 and E2F2 induction in response to DNA damage preserves genomic stability in neuronal cells. Cell Cycle 14, 1300–1314. 10.4161/15384101.2014.985031.

37. Chen, H.R., Lin, G.T., Huang, C.K., and Fann, M.J. (2014). Cdk12 and Cdk13 regulate axonal elongation through a common signaling pathway that modulates Cdk5 expression. Exp Neurol 261, 10–21. 10.1016/j.expneurol.2014.06.024.

38. Cortés, N., Guzmán-Martínez, L., Andrade, V., González, A., and Maccioni, R.B. (2019). CDK5: A Unique CDK and Its Multiple Roles in the Nervous System. J Alzheimers Dis 68, 843–855. 10.3233/Jad-180792.

39. Upadhyay, A., Joshi, V., Amanullah, A., Mishra, R., Arora, N., Prasad, A., and Mishra, A. (2017). E3 Ubiquitin Ligases Neurobiological Mechanisms: Development to Degeneration. Front Mol Neurosci 10. ARTN 15110.3389/fnmol.2017.00151.

40. Simoes, A.R., and Rhiner, C. (2017). A Cold-Blooded View on Adult Neurogenesis. Front Neurosci-Switz 11. ARTN 32710.3389/fnins.2017.00327.

41. Zars, T., Fischer, M., Schulz, R., and Heisenberg, M. (2000). Localization of a Short-Term Memory in Drosophila. Science 288, 672–675. 10.1126/science.288.5466.672.

42. Lee, T., and Luo, L. (2001). Mosaic analysis with a repressible cell marker (MARCM) for Drosophila neural development. Trends Neurosci 24, 251–254. 10.1016/s0166-2236(00)01791-4.

43. Gelbart, W.M. (1992). The Genome of Drosophila-Melanogaster - Lindsley,Dl, Zimm,Gg. Science 257, 1421–1422. DOI 10.1126/science.257.5075.1421.

44. Ward, L. (1923). The Genetics of Curly Wing in Drosophila. Another Case of Balanced Lethal Factors. Genetics 8, 276–300. 10.1093/genetics/8.3.276.

45. Kraut, R., and CamposOrtega, J.A. (1996). inscuteable, a neural precursor gene of Drosophila, encodes a candidate for a cytoskeleton adaptor protein. Dev Biol 174, 65–81. DOI 10.1006/dbio.1996.0052.

46. Albertson, R., Chabu, C., Sheehan, A., and Doe, C.Q. (2004). Scribble protein domain mapping reveals a multistep localization mechanism and domains necessary for establishing cortical polarity. J Cell Sci 117, 6061–6070. 10.1242/jcs.01525.

47. Yang, C.P., Samuels, T.J., Huang, Y., Yang, L., Ish-Horowicz, D., Davis, I., and Lee, T. (2017). Imp and Syp RNA-binding proteins govern decommissioning of Drosophila neural stem cells. Development 144, 3454–3464. 10.1242/dev.149500.

48. Pahl, M.C., Doyle, S.E., and Siegrist, S.E. (2019). E93 Integrates Neuroblast Intrinsic State with Developmental Time to Terminate MB Neurogenesis via Autophagy. Curr Biol 29, 750–762.e753. 10.1016/j.cub.2019.01.039.

49. Connolly, J.B., Roberts, I.J.H., Armstrong, J.D., Kaiser, K., Forte, M., Tully, T., and O’Kane, C.J. (1996). Associative Learning Disrupted by Impaired Gs Signaling in Drosophila Mushroom Bodies. Science 274, 2104–2107. 10.1126/science.274.5295.2104.

50. Bahrampour, S., Gunnar, E., Jonsson, C., Ekman, H., and Thor, S. (2017). Neural Lineage Progression Controlled by a Temporal Proliferation Program. Dev Cell 43, 332–348.e334. 10.1016/j.devcel.2017.10.004.

51. Caygill, E.E., and Brand, A.H. (2016). The GAL4 System: A Versatile System for the Manipulation and Analysis of Gene Expression. Methods Mol Biol 1478, 33–52. 10.1007/978-1-4939-6371-3_2.

52. Wieschaus, E., Nusslein-Volhard, C., and Kluding, H. (1984). Krüppel, a gene whose activity is required early in the zygotic genome for normal embryonic segmentation. Dev Biol 104, 172–186. 10.1016/0012-1606(84)90046-0.

53. Knipple, D.C., Seifert, E., Rosenberg, U.B., Preiss, A., and Jäckle, H. (1985). Spatial and temporal patterns of Krüppel gene expression in early Drosophila embryos. Nature 317, 40–44. 10.1038/317040a0.

54. Jacob, J., Maurange, C., and Gould, A.P. (2008). Temporal control of neuronal diversity: common regulatory principles in insects and vertebrates? Development 135, 3481–3489. 10.1242/dev.016931.

55. Li, X., Chen, Z.Q., and Desplan, C. (2013). Temporal Patterning of Neural Progenitors in Drosophila. Curr Top Dev Biol 105, 69–96. 10.1016/B978-0-12-396968-2.00003-8.

56. Dubuis, J.O., Samanta, R., and Gregor, T. (2013). Accurate measurements of dynamics and reproducibility in small genetic networks. Molecular Systems Biology 9, 639. 10.1038/msb.2012.72.

57. Kosman, D., Small, S., and Reinitz, J. (1998). Rapid preparation of a panel of polyclonal antibodies to Drosophila segmentation proteins. Development Genes and Evolution 208, 290–294. 10.1007/s004270050184.

58. Kunz, T., Kraft, K.F., Technau, G.M., and Urbach, R. (2012). Origin of Drosophila mushroom body neuroblasts and generation of divergent embryonic lineages. Development 139, 2510–2522. 10.1242/dev.077883.

59. Venken, K.J.T., Carlson, J.W., Schulze, K.L., Pan, H., He, Y., Spokony, R., Wan, K.H., Koriabine, M., de Jong, P.J., White, K.P., et al. (2009). Versatile P[acman] BAC libraries for transgenesis studies in Drosophila melanogaster. Nature Methods 6, 431–434. 10.1038/nmeth.1331.

60. Liu, Z., Yang, C.-P., Sugino, K., Fu, C.-C., Liu, L.-Y., Yao, X., Lee, L.P., and Lee, T. (2015). Opposing intrinsic temporal gradients guide neural stem cell production of varied neuronal fates. Science 350, 317–320. 10.1126/science.aad1886.

61. Kang, P., Chang, K., Liu, Y., Bouska, M., Birnbaum, A., Karashchuk, G., Thakore, R., Zheng, W., Post, S., Brent, C.S., et al. (2017). Drosophila Kruppel homolog 1 represses lipolysis through interaction with dFOXO. Scientific Reports 7, 16369. 10.1038/s41598-017-16638-1.

62. Islam, I.M., and Erclik, T. (2022). Imp and Syp mediated temporal patterning of neural stem cells in the developing Drosophila CNS. Genetics 222, iyac103. 10.1093/genetics/iyac103.

63. Stratmann, J., Gabilondo, H., Benito-Sipos, J., and Thor, S. (2016). Neuronal cell fate diversification controlled by sub-temporal action of Kruppel. Elife 5. 10.7554/eLife.19311.

64. Kraft, K.F., Massey, E.M., Kolb, D., Walldorf, U., and Urbach, R. (2016). Retinal homeobox promotes cell growth, proliferation and survival of mushroom body neuroblasts in the Drosophila brain. Mechanisms of Development 142, 50–61. 10.1016/j.mod.2016.07.003.

65. Zhu, S., Lin, S., Kao, C.F., Awasaki, T., Chiang, A.S., and Lee, T. (2006). Gradients of the Drosophila Chinmo BTB-zinc finger protein govern neuronal temporal identity. Cell 127, 409–422. 10.1016/j.cell.2006.08.045.

66. Maurange, C., Cheng, L., and Gould, A.P. (2008). Temporal transcription factors and their targets schedule the end of neural proliferation in. Cell 133, 891–902. 10.1016/j.cell.2008.03.034.

67. Cayre, M., Strambi, C., Charpin, P., Augier, R., Meyer, M.R., Edwards, J.S., and Strambi, A. (1996). Neurogenesis in adult insect mushroom bodies. J Comp Neurol 371, 300–310. 10.1002/(SICI)1096-9861(19960722)371:2<300::AID-CNE9>3.0.CO;2-6.

68. Farris, S.M., and Sinakevitch, I. (2003). Development and evolution of the insect mushroom bodies: towards the understanding of conserved developmental mechanisms in a higher brain center. Arthropod Struct Dev 32, 79–101. 10.1016/s1467-8039(03)00009-4.

69. Takahashi, K., and Yamanaka, S. (2006). Induction of pluripotent stem cells from mouse embryonic and adult fibroblast cultures by defined factors. Cell 126, 663–676. DOI 10.1016/j.cell.2006.07.024.

70. Han, M.J., Lee, W.J., Choi, J., Hong, Y.J., Uhm, S.J., Choi, Y., and Do, J.T. (2021). Inhibition of neural stem cell aging through the transient induction of reprogramming factors. J Comp Neurol 529, 595–604. 10.1002/cne.24967.

71. McAvoy, K.M., Scobie, K.N., Berger, S., Russo, C., Guo, N.N., Decharatanachart, P., Vega-Ramirez, H., Miake-Lye, S., Whalen, M., Nelson, M., et al. (2016). Modulating Neuronal Competition Dynamics in the Dentate Gyrus to Rejuvenate Aging Memory Circuits. Neuron 91, 1356–1373. 10.1016/j.neuron.2016.08.009.

72. Guo, N.N., McDermott, K.D., Shih, Y.T., Zanga, H., Ghosh, D., Herber, C., Meara, W.R., Coleman, J., Zagouras, A., Wong, L.P., et al. (2022). Transcriptional regulation of neural stem cell expansion in the adult hippocampus. Elife 11. ARTN e7219510.7554/eLife.72195.

73. Kimata, Y., Martins, T., Meghini, F., and Florio, F. (2016). The APC/C Coordinates Retinal Differentiation with G1 Arrest through the Nek2-Dependent Modulation of Wingless Signaling. Dev Cell 40. 10.1016/j.devcel.2016.12.005.

74. Wu, J.S., and Luo, L.Q. (2006). A protocol for dissecting brains for live imaging or immunostaining. Nat Protoc 1, 2110–2115. 10.1038/nprot.2006.336.

75. Schneider, C.A., Rasband, W.S., and Eliceiri, K.W. (2012). NIH Image to ImageJ: 25 years of image analysis. Nat Methods 9, 671–675. 10.1038/nmeth.2089.

76. Stuart, T., Butler, A., Hoffman, P., Hafemeister, C., Papalexi, E., Mauck, W.M., Hao, Y.H., Stoeckius, M., Smibert, P., and Satija, R. (2019). Comprehensive Integration of Single-Cell Data. Cell 177, 1888-+. 10.1016/j.cell.2019.05.031.

