## Supplementary materials for "The Conserved Transcription Factor Krüppel Regulates the Survival and Neurogenesis of Mushroom Body Neuroblasts in *Drosophila* Adult Brains"

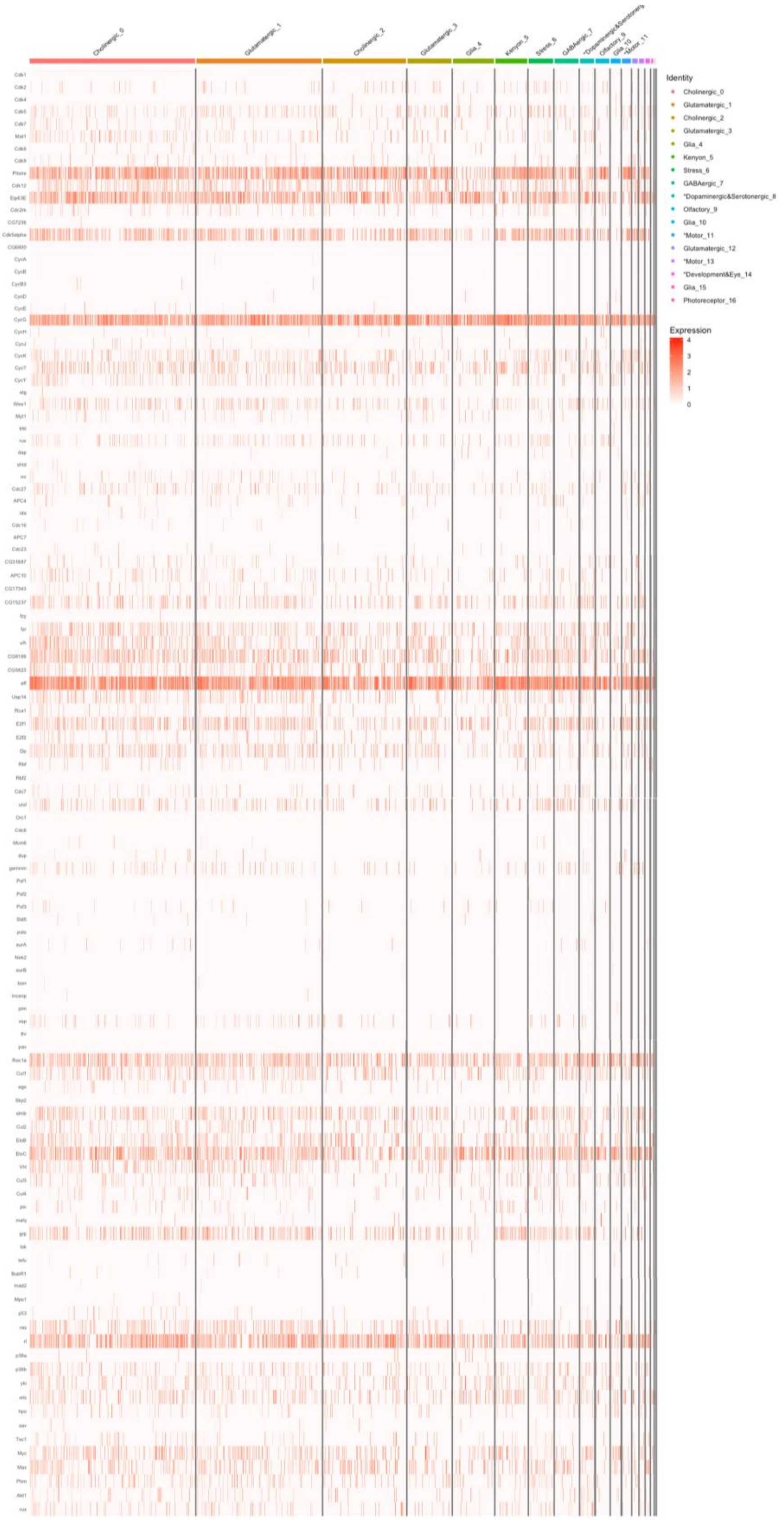


**Figure S1. Heatmap of CCR gene expression in 17 clusters of *Drosophila* adult brain**

The heatmap displays the expression levels of 112 selected CCR genes across 17 identified clusters in the Drosophila adult brain. Each row represents a CCR gene, and each column corresponds to an individual cell within the clusters. The colour intensity indicates the expression level, with darker red indicating higher expression levels and lighter shades indicating lower expression levels. This visualization highlights the heterogeneity of CCR gene expression within each cluster.

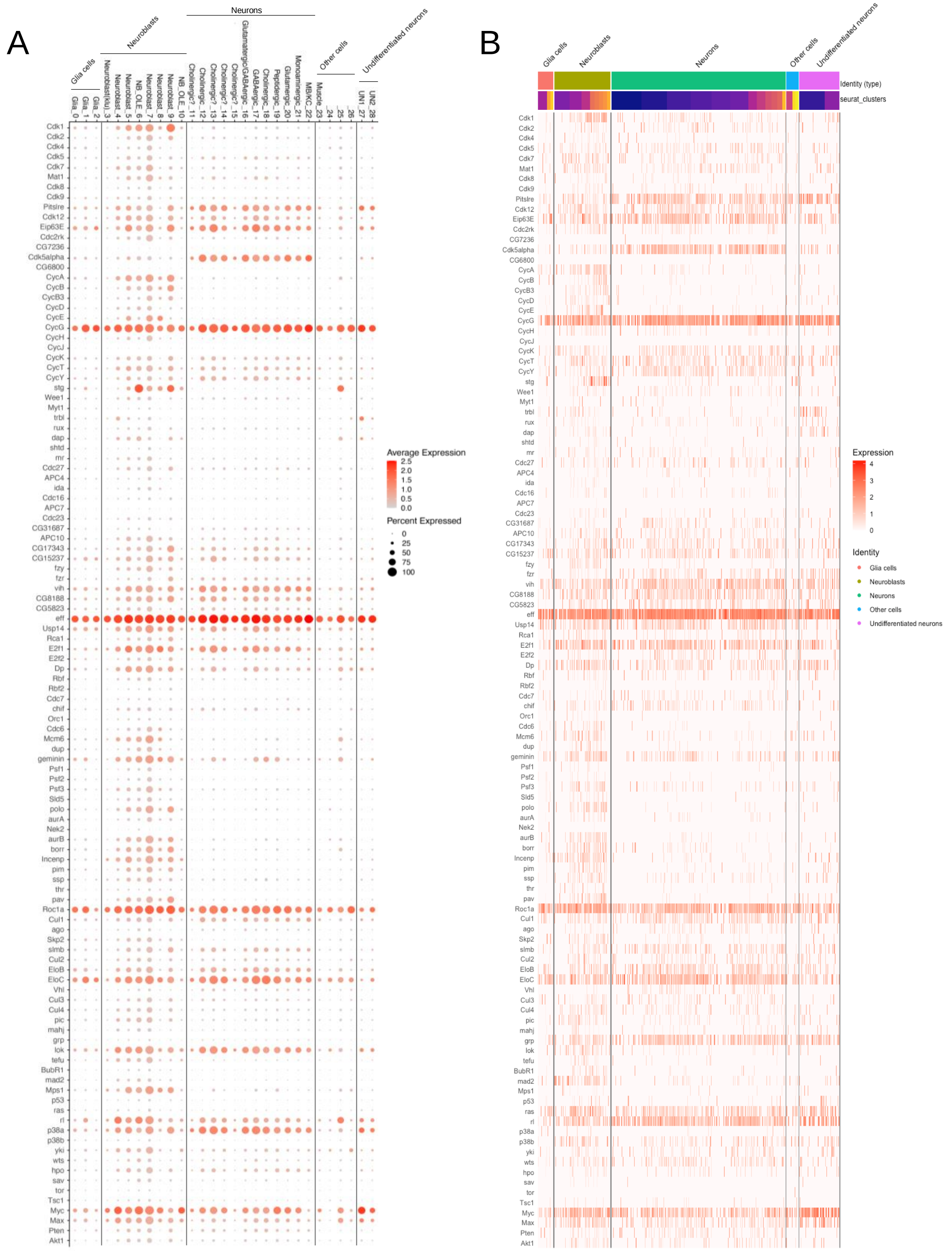


**Figure S2. Dot plot and heatmap of CCR gene expression across clusters of the *Drosophila* larval brain.** (A) Dot plot showing the expression patterns of CCR genes across 29 cell clusters identified through Seurat using the recent single-cell RNA sequencing data from *Drosophila* larval brains *(Avalos et al., 2019).* The colour intensity of each dot represents the average expression level of the CCR gene, with darker red indicating higher expression levels similar to Fig. 1D. The size of each dot indicates the proportion of cells within the cluster expressing the gene, with larger dots representing higher proportions.) and dot size representing the proportion of cells expressing each gene. (B) Heatmap displaying the expression levels of CCR genes in *Drosophila* larval brains, presented similarly to Fig. S1. Each row represents a CCR gene, and each column corresponds to an individual cell, with colour intensity indicating the expression level.

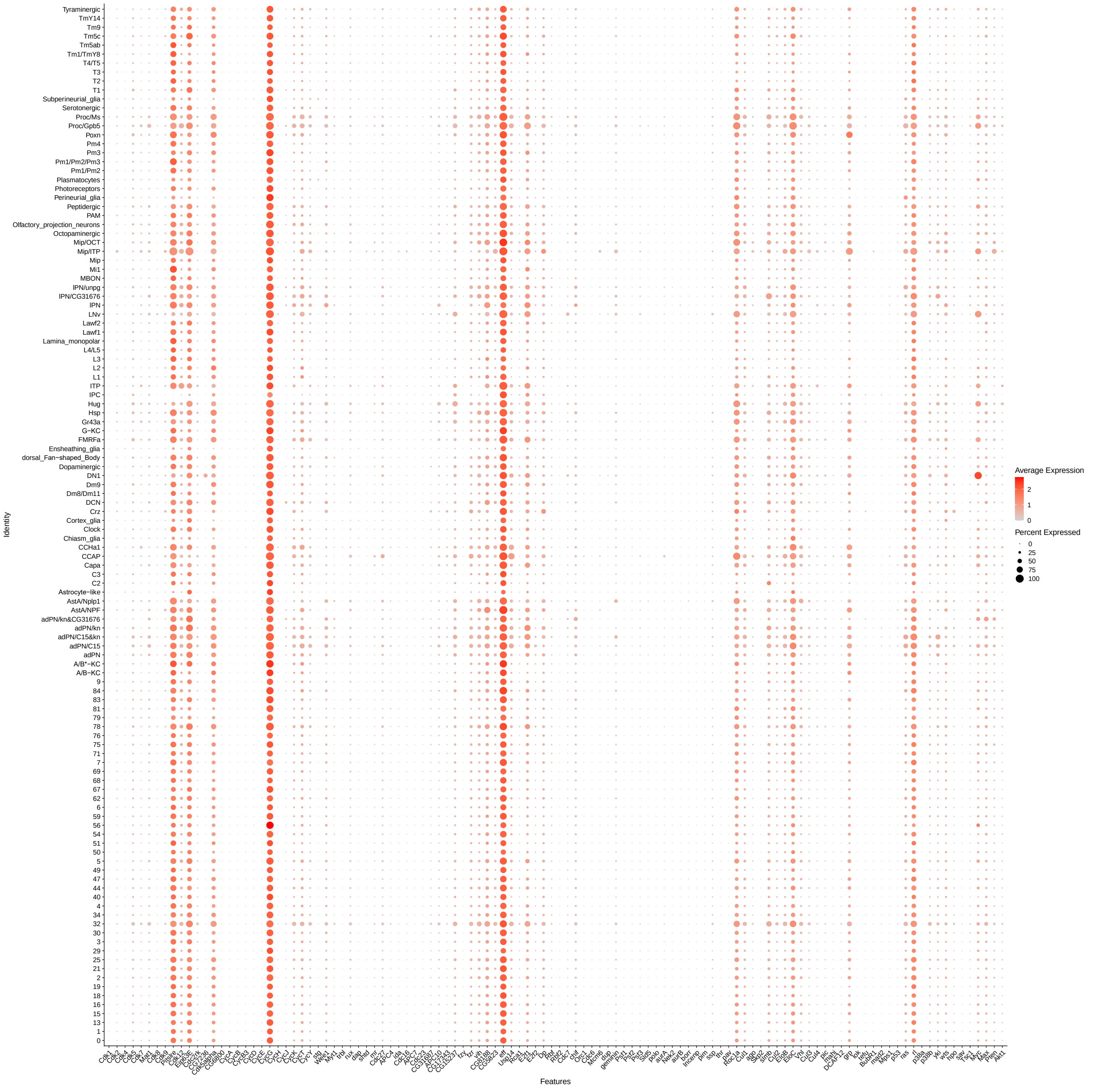


**Figure S3 Dot plot of CCR gene expression across the 84 clusters of the *Drosophila* adult brain, identified in** *Davie, Janssens et al. 2018*.

The dot plot illustrates the expression of 112 selected CCR genes across 84 identified clusters in the Drosophila adult brain. Each dot represents the average expression level of a CCR gene within a cluster, with the colour intensity indicating the expression level (darker red indicates higher expression). The size of each dot represents the proportion of cells within the cluster expressing the gene, with larger dots indicating higher proportions. This plot provides an overview of the distribution and prevalence of CCR gene expression across all identified clusters.

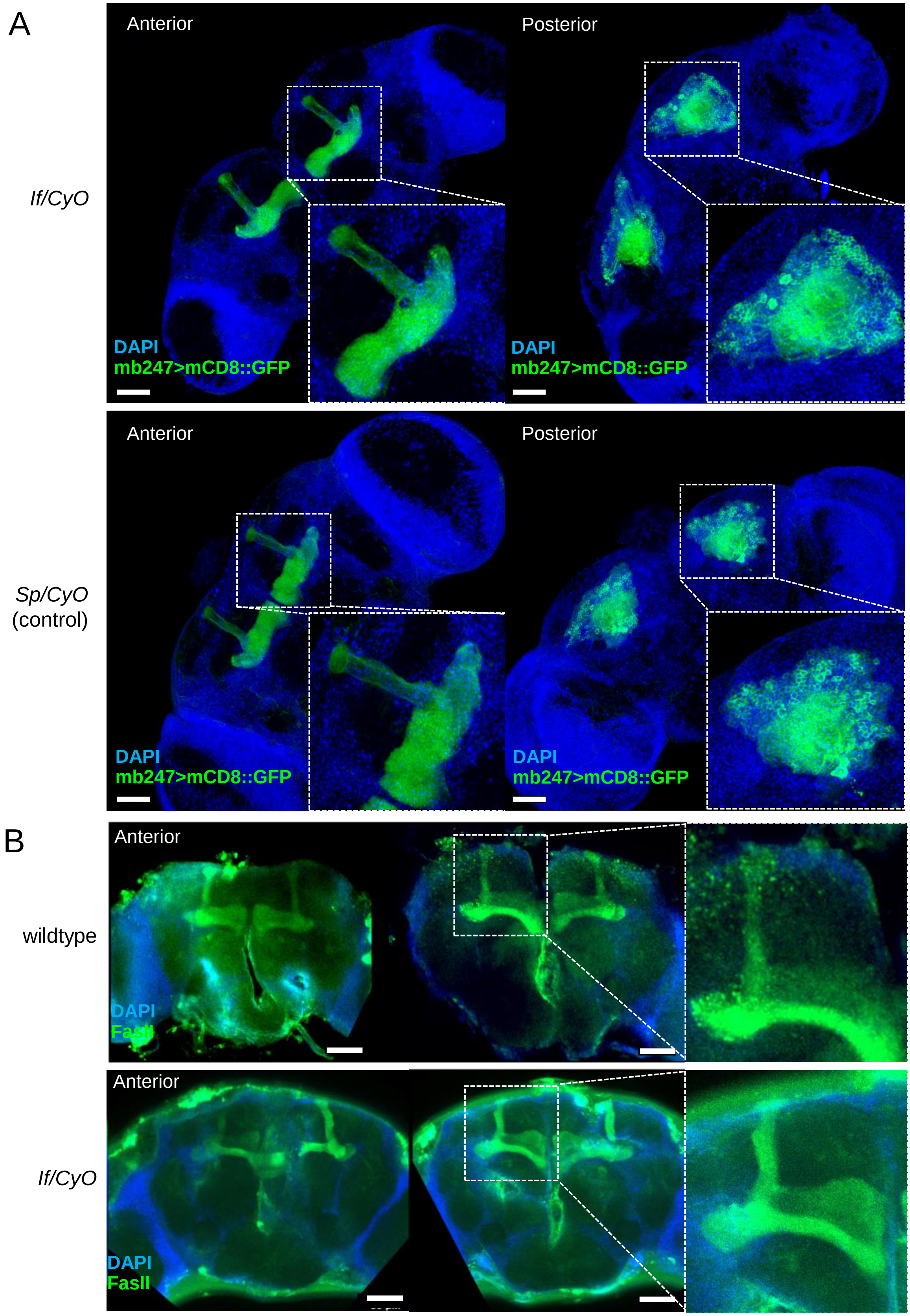


**Figure S4 The overall structures of the adult mushroom bodies are not altered in *If* mutants**
(A) The structures of the MBs were visualised by *mb247>mCD8::GFP* in *If* mutant (*If/CyO,* top) and control (*Sp/CyO*, bottom) adult brains. The anterior and posterior views of the adult brains, where MB lobes, calyx, and cell bodies are visible, are shown on the left and right, respectively. DAPI (blue) and GFP (green). Insets show magnified images. Scale bars: 50 µm. (B) Anterior views of the 3D-reconstructed confocal images of the central brains in wild type (top) and the *If* mutant (*If/CyO,* bottom) where MB lobes were visualised using anti-FasII antibody (green). No obvious morphological alterations were observed. Scale bars: 50 µm.

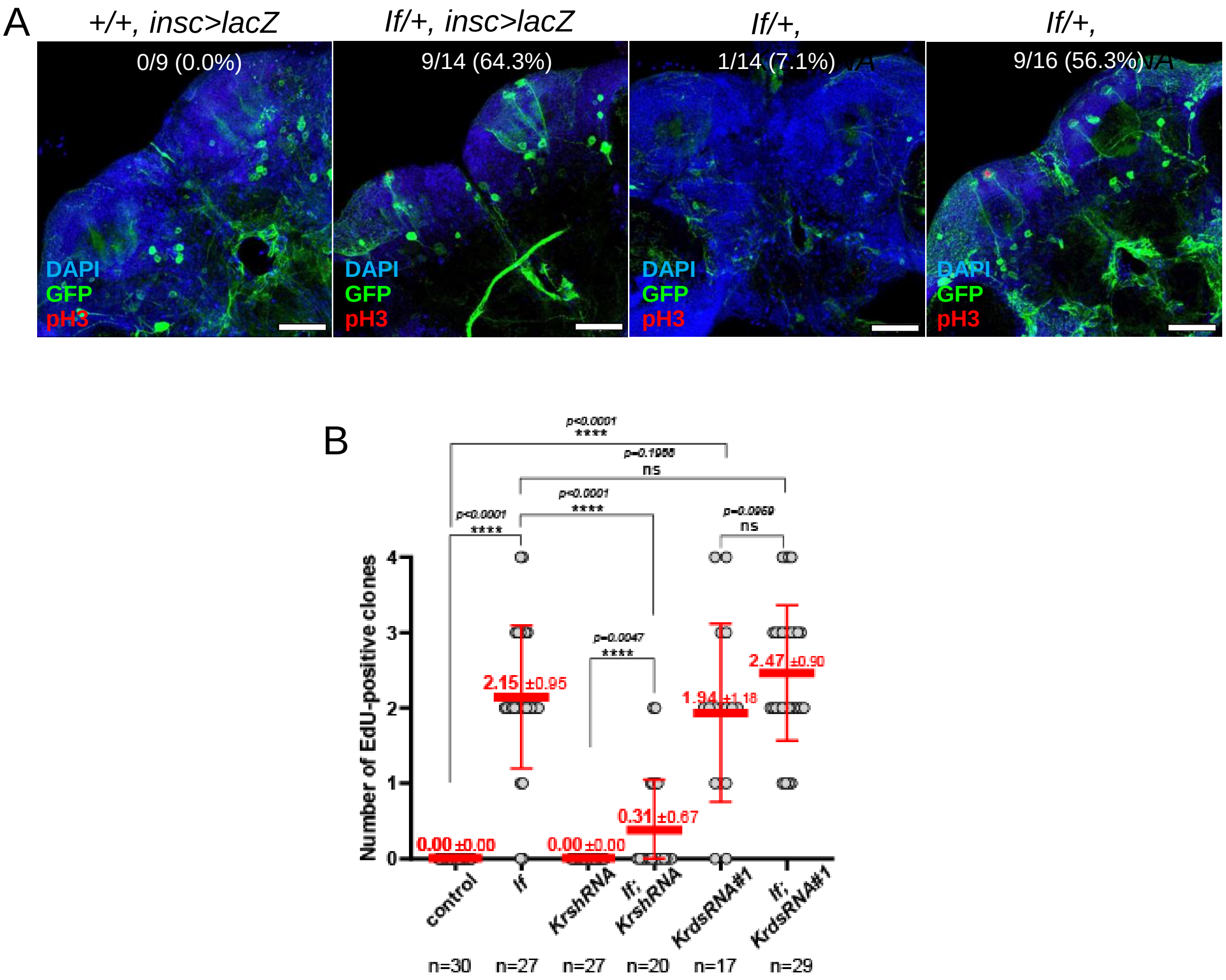


**Figure S5 Kr knockdown using *Kr*shRNA induction suppresses ectopic NB formation in *If* mutant adult brains.**

(A) Representative images of adult central brains from control (*+/+, insc>lacZ*), *If* mutant (*If/+, insc>lacZ*), *If* mutant with NB-specific *KrshRNA* induction (*If/+, insc>KrshRNA*), and *If* mutant with NB-specific *HbshRNA* induction (*If/+, insc-gal4>HbshRNA*). DNA is shown in blue, *insc*-gal4-driven mCD8::GFP in green, and pH3 in red. The numbers and ratios of brains exhibiting pH3-positive cells among the total number of brains examined are quantified and shown at the top. Scale bars: 50 µm. Mitotic NB formation in *If* mutant adult brains was partially suppressed by NB-specific induction of *KrshRNA*, but not *HbshRNA*. (B) EdU labelling was conducted on adult flies of the indicated strains for three to five days post-eclosion. Brains were subsequently dissected, and the number of EdU-positive cell clones per hemisphere was quantified and displayed in scattered dot plots. Means and standard deviations are represented by thick and thin red bars, respectively, with actual values displayed above the bars in red. 'n' denotes the number of brains analysed per condition. Statistical significance was determined using the Mann-Whitney U test, with significance levels denoted as: * ($p \leq 0.05$), ** ($p \leq 0.01$), *** ($p \leq 0.001$), **** ($p < 0.0001$). 'ns' indicates non-significant results.

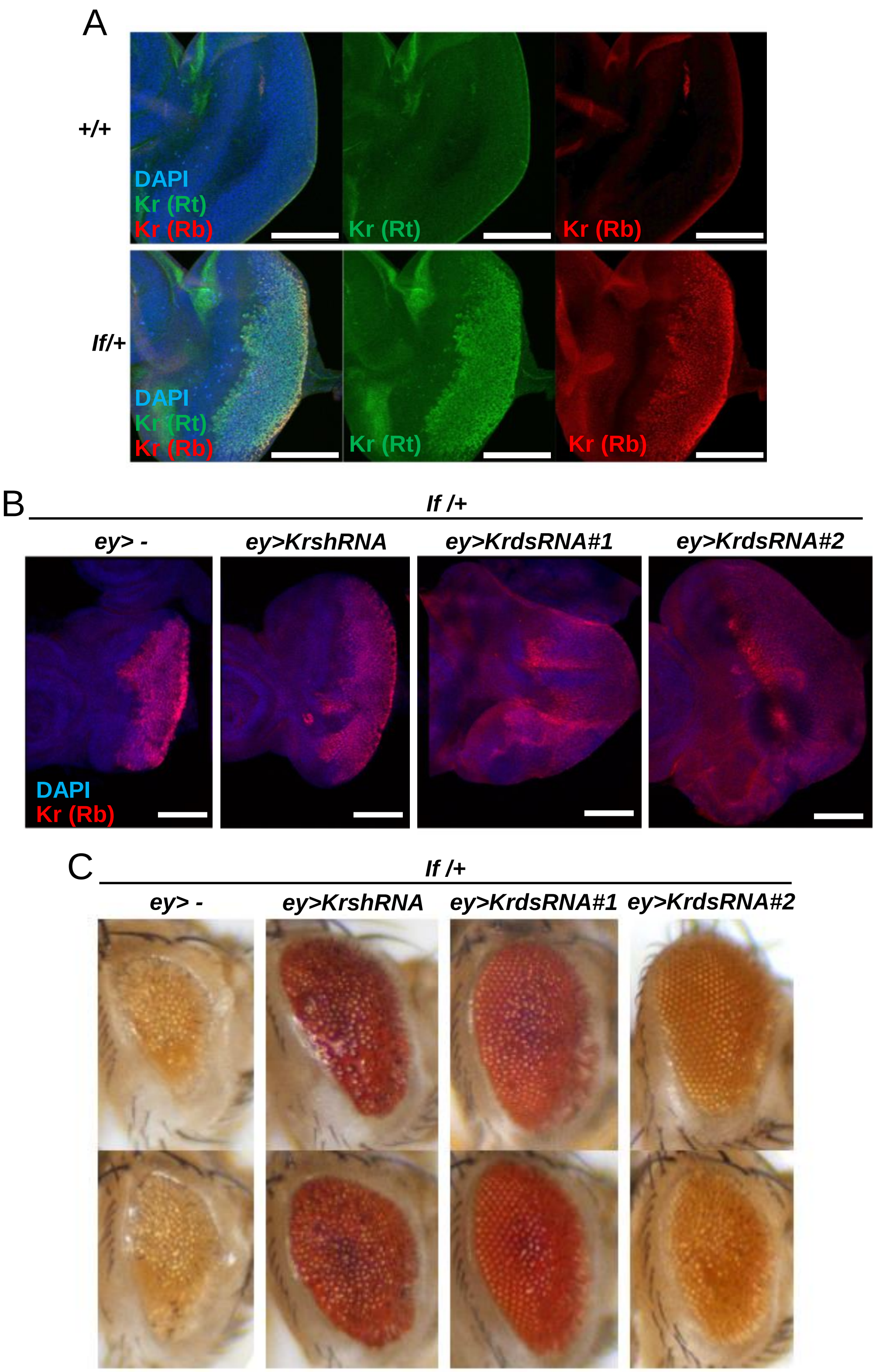


**Figure S6 The *KrdsRNA* constructs repress Kr expression more efficiently than the *Kr*shRNA construct** (A) Eye imaginal discs from of wild-type (+/+) and *If* mutant (*If*/+) third instar larvae stained by DAPI (blue), anti-Kr antibodies raised in a rat (Rt, green) and a rabbit (Rb, red)(Kosman, Small et al. 1998)(Dubuis, Samanta et al. 2013). Overexpression of Kr proteins was detected by both anti-Kr antibodies in differentiating cells in *If* mutant eye discs, but not in wild-type discs. Scale bars: 100 µm. (B) Eye imaginal discs of *If* mutants, either uninduced (-) or expressing *KrshRNA*, *KrdsRNA#1*, or *KrdsRNA#2* driven by the eye region-specific *eyeless*-gal4 (*ey*-gal4). Kr protein levels were detected using the rabbit anti-Kr antibody, verified for specificity and linearity previously(Dubuis, Samanta et al. 2013). DNA is shown in blue, and Kr signals in red. Scale bars: 100 µm. (C) Representative images of adult eyes of *If* mutants with no UAS construct, or *KrshRNA*, *KrdsRNA#1*, or *KrdsRNA#2* induced by *ey*-gal4. The *Kr*dsRNA constructs more effectively suppressed ectopic Kr expression in the eye imaginal discs of If mutants and more efficiently ameliorated morphological defects in the adult compound eyes than the *Kr*shRNA construct.

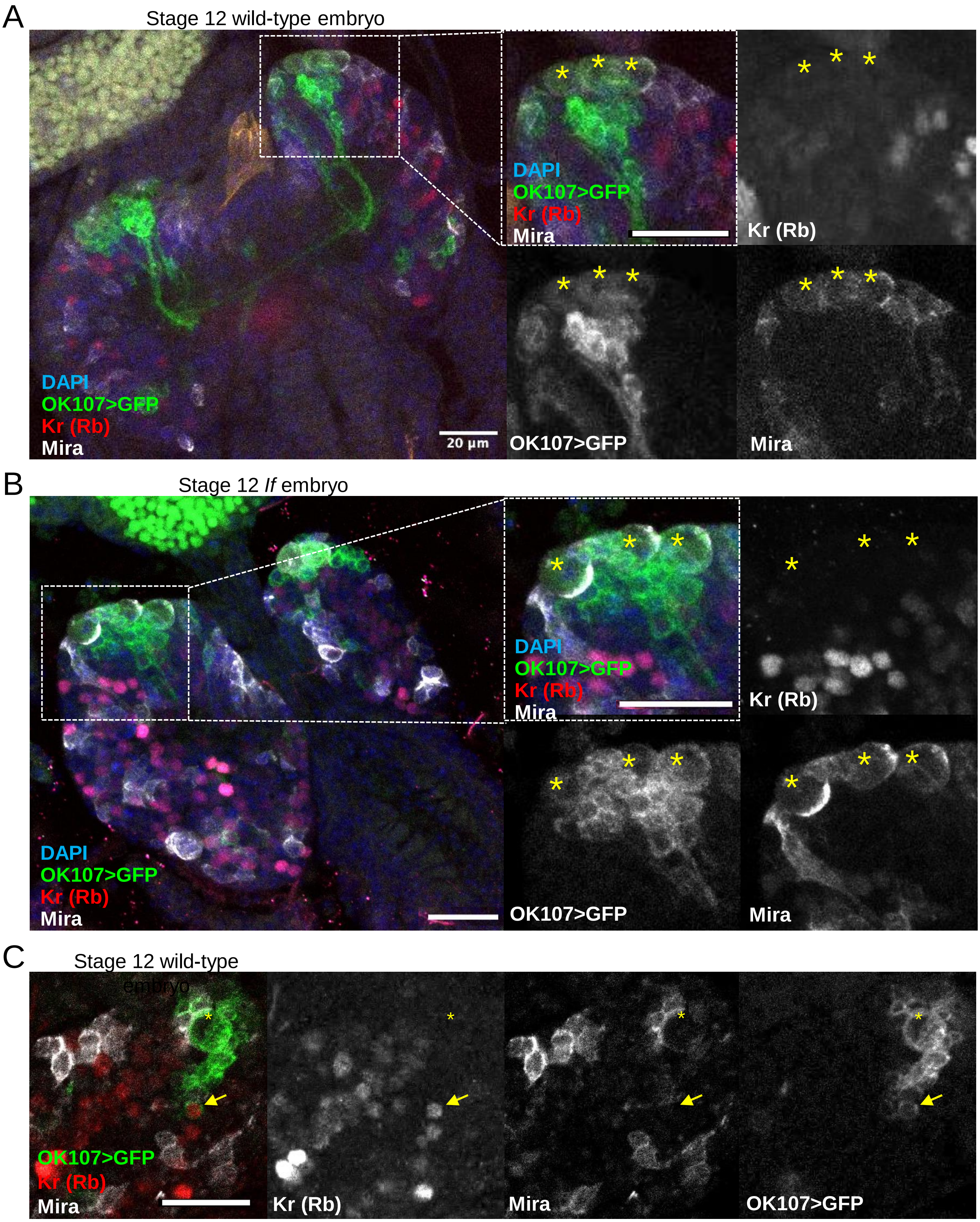


**Figure S7 Kr is undetectable in MBNBs in embryos**

(A, B) Representative images of the CNS in wild-type Stage 12 embryos. Kr proteins were detected using the rabbit anti-Kr antibody(Dubuis, Samanta et al. 2013). In merged images, DNA is shown in blue, *OK107*-gal4-driven mCD8::GFP in green, Kr in red, and Mira in grey, with accompanying mono-coloured images for each signal in grey. Scale bars: 20 µm. Asterisks indicate MBNBs, marked by the co-expression of *OK107*>GFP and Mira. (B) Arrows point to a neuronal cell in the MBNB lineage that shows clear Kr signals. While Kr is expressed in various cells within the embryonic CNS, it appears absent in MBNBs.

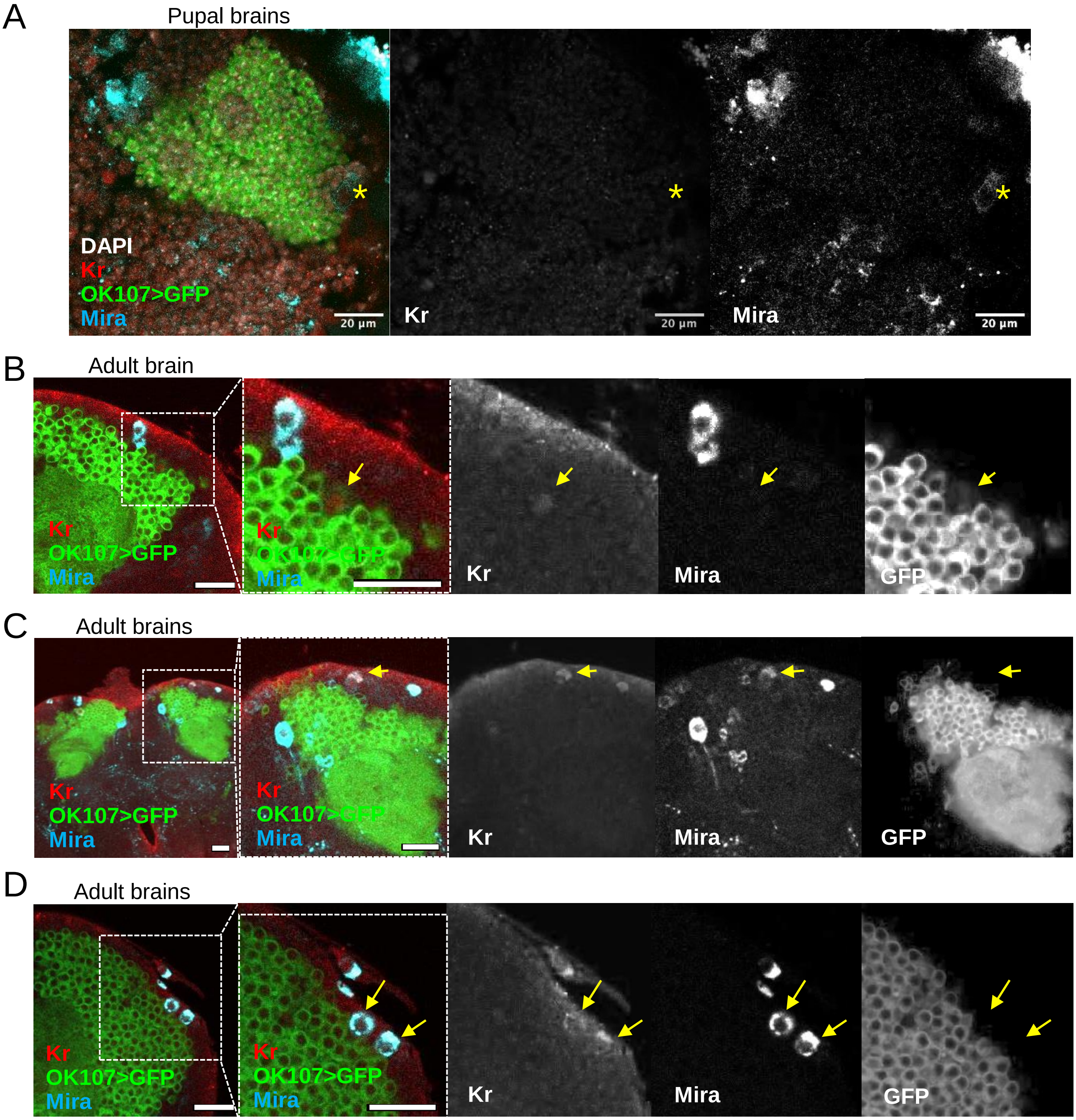


**Figure S8 Kr is undetectable in MBNBs in pupal and adult brains**
(A) Representative images of the dorsoposterior MB cell body region in pupal brains. In merged images, Kr-specific signals, detected using a rabbit anti-Kr antibody, are shown in red, DNA in grey, OK107>GFP in green, and Mira in cyan. Mono-coloured views are provided for Kr and Mira. Asterisks indicate MBNBs where Kr signals were not detectable. (B-D) Images of the dorsoposterior region of the wild-type adult brain, focusing on MB cell bodies and the Calyx. In merged images, Kr signals are shown in red, OK107-driven GFP in green, and Mira in cyan, with single-coloured views for each signal provided alongside the merged images. No MBNBs were detected in wild-type adult brains. (B) Shows a MB neuron with weak nuclear Kr signals (arrow). (C, D) Display uncharacterised cells near the MB cell body, exhibiting cytoplasmic Kr signals co-localised with Mira (arrows). Scale bars: 20 µm

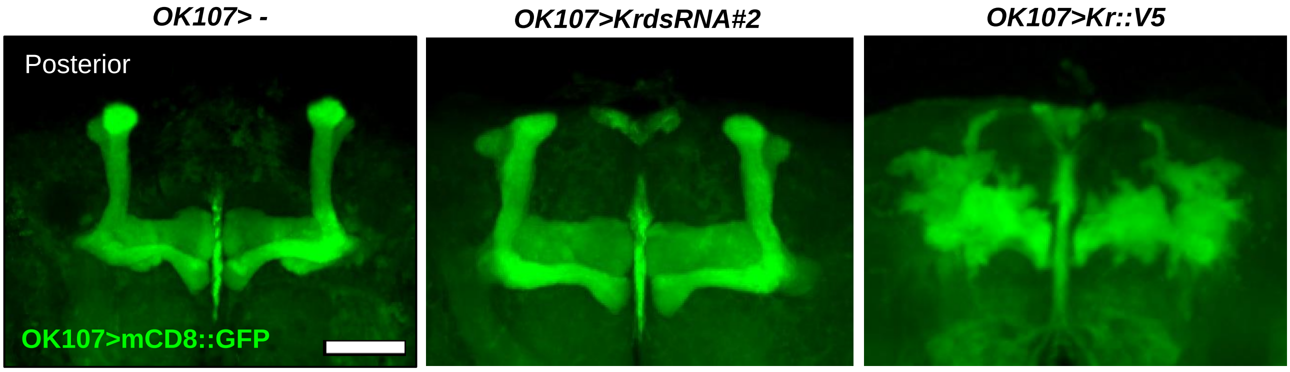


**Figure S9 Effects of Kr depletion or overexpression in MB lineages using *OK107*-gal4**
Posterior views of the 3D-reconstructed confocal images of the central brains of the control (no UAS gene), *OK107>KrdsRNA#2,* and *OK107>Kr::V5* adult flies where MBs were visualised by mCD8::GFP induced by *OK107*-gal4 (green). While the control and *OK107>KrdsRNA#2* adult brains did not exhibit any defects, Kr::V5 overexpression driven by *OK107*-gal4 caused high lethality rate and surviving adult flies displayed severe morphological defects in the MB structure. Scale bars: 50 µm.

### References


1. Brunet Avalos, C., G. L. Maier, R. Bruggmann and S. G. Sprecher (2019). "Single cell transcriptome atlas of the Drosophila larval brain." Elife **8**.

2. *Kosman, D., S. Small and J. Reinitz (1998). "Rapid preparation of a panel of polyclonal antibodies to Drosophila segmentation proteins." Development Genes and Evolution* ***208****(5): 290-294.*

3. *Dubuis, J. O., R. Samanta and T. Gregor (2013). "Accurate measurements of dynamics and reproducibility in small genetic networks." Molecular Systems Biology* ***9****(1): 639.*

4. *Davie, K., J. Janssens, D. Koldere, M. De Waegeneer, U. Pech, L. Kreft, S. Aibar, S. Makhzami, V. Christiaens, C. B. González-Blas, S. Poovathingal, G. Hulselmans, K. I. Spanier, T. Moerman, B. Vanspauwen, S. Geurs, T. Voet, J. Lammertyn, B. Thienpont, S. Liu, N. Konstantinides, M. Fiers, P. Verstreken and S. Aerts (2018). "A Single-Cell Transcriptome Atlas of the Aging Drosophila Brain." Cell 174(4): 982-+.*
